# OGT prevents DNA demethylation and suppresses the expression of transposable elements in heterochromatin by restraining TET activity genome-wide

**DOI:** 10.1101/2024.01.31.578097

**Authors:** Hugo Sepulveda, Xiang Li, Xiaojing Yue, J. Carlos Angel, Leo J. Arteaga-Vazquez, Caitlin Brown, Melina Brunelli, Natasha Jansz, Fabio Puddu, Jamie Scotcher, Páidí Creed, Patrick Kennedy, Cindy Manriquez, Samuel A Myers, Robert Crawford, Geoffrey J. Faulkner, Anjana Rao

**Author notes:** These authors contributed equally (Hugo Sepulveda, Xiang Li).

## Abstract

The *O-*GlcNAc transferase OGT interacts robustly with all three mammalian TET methylcytosine dioxygenases. We show here that deletion of the *Ogt* gene in mouse embryonic stem cells (mESC) results in a widespread increase in the TET product 5-hydroxymethylcytosine (5hmC) in both euchromatic and heterochromatic compartments, with concomitant reduction of the TET substrate 5-methylcytosine (5mC) at the same genomic regions. mESC engineered to abolish the TET1-OGT interaction likewise displayed a genome-wide decrease of 5mC. DNA hypomethylation in OGT-deficient cells was accompanied by de-repression of transposable elements (TEs) predominantly located in heterochromatin, and this increase in TE expression was sometimes accompanied by increased *cis*-expression of genes and exons located 3’ of the expressed TE. Thus, the TET-OGT interaction prevents DNA demethylation and TE expression in heterochromatin by restraining TET activity genome-wide. We suggest that OGT protects the genome against DNA hypomethylation and impaired heterochromatin integrity, preventing the aberrant increase in TE expression observed in cancer, autoimmune-inflammatory diseases, cellular senescence and ageing.

## INTRODUCTION

DNA cytosine methylation and demethylation are controlled by DNA methyltransferases (DNMTs) and TET (<u>T</u>en-<u>E</u>leven <u>T</u>ranslocation) methylcytosine dioxygenases, respectively [1–7]. The three mammalian members of the TET family catalyze the oxidation of 5-methylcytosine (5mC) to 5-hydroxymethylcytosine (5hmC) and beyond [8–11]. Together, the oxidized methylcytosines are intermediates in both the “passive” replication-dependent and the “active” replication-independent pathways of DNA demethylation (reviewed in [5–7]).

OGT (*O-*GlcNAc transferase) is an essential X-chromosome-encoded enzyme that modifies the hydroxyl groups of Ser/Thr residues in many intracellular (cytoplasmic and nuclear) proteins with N-acetylglucosamine (GlcNAc) [12–18]. The *O-*GlcNAc modification is reversed by O-GlcNAcase (OGA) [12]. OGT is essential for the very earliest stages of embryonic development in mammals: deletion of the *Ogt* gene in the mouse germline results in embryonic lethality prior to blastocyst implantation [19, 20]. In all primary cells and cell lines examined, OGT is essential for proliferation and cell survival [12, 21].

OGT binds tightly to all three mammalian TET proteins, TET1, TET2 and TET3 [22–24]. TET proteins recruit OGT to chromatin [22], and all three TET enzymes are *O*-GlcNAcylated by OGT [21, 22, 24–28], but the functional consequences of the TET-OGT interaction are not well understood. Deletion of the gene encoding the highly *O*-GlcNAcylated scaffold protein PROSER1 [21, 28] in HEK293T cells resulted in a partial decrease of *O*-GlcNAcylation of TET2 proteins and reduced TET2 protein levels [29]; however, *PROSER1* deletion was accompanied by altered methylation of only a small fraction of genomic regions [29], prompting us to investigate the role of the upstream regulator OGT in globally modulating TET function and DNA modifications in the genome.

A major function of DNA methylation is the transcriptional repression of transposable elements (TEs), imprinted genes, satellite repeats and other repetitive elements ([30–34]; reviewed in [35–38]). TEs include long terminal repeat (LTR)-containing elements – endogenous retroviruses (ERVs) and intracisternal A-type particles (IAPs) – as well as non-LTR retrotransposons (long and short interspersed elements, LINEs and SINEs) [36–39]. The expression of TEs, especially ERVs and all but the youngest LINE-1 (L1) families, is suppressed by the transcriptional co-repressor KAP1 (also known as TRIM28), which is recruited to TE sequences by an extensive family of KRAB zinc finger proteins that co-evolved with TEs in an ‘arms race’ to suppress TE expression [33, 34, 36–44]. The functions of TETs, DNMTs and the TET-OGT interaction have also been explored in the context of TE expression [45, 46]. In mouse embryonic stem cells (mESC), TET1 and TET2 bound the regulatory regions of diverse classes of TEs, including young L1 subfamilies that are not yet repressed by KAP1; TET binding at these elements was associated with increased 5hmC and decreased 5mC [45], implicating TET-mediated DNA demethylation in TE expression. In “naïve” mESC cultured under 2i conditions, which display lower 5mC levels and higher TE expression than “primed” mESC grown in serum plus LIF [47, 48], depletion of the H3K9 methyltransferase SETDB1 increased the expression of IAPs and young L1 (T_F_ subfamily) elements, and this increase required TET catalytic function [46]. KAP1 and TET proteins are implicated in controlling TE expression via H3K9 and DNA methylation [49].

In this study, we explored the connection between OGT and DNA methylation/demethylation pathways using *Ogt fl*, *Cre-ERT2* mESC that could be inducibly deleted for the *Ogt* gene by treatment with 4-hydroxytamoxifen (4-OHT) [21]. Using a recently developed six-letter sequencing method that simultaneously detects 5hmC and 5mC (as well as all four canonical bases) in a single sequencing run [50], we report that acute deletion of the *Ogt* gene results in a global loss of 5mC and a global increase of 5hmC that occurs in both euchromatic and heterochromatic compartments within 6 days of 4-OHT exposure. Loss of 5mC was accompanied by decreased protein levels of DNMT1 and its partner UHRF1; increased expression of TEs – the LINE-1 (L1) family and the MERVL (MT2) and IAP subfamilies – that are predominantly located in heterochromatin; and dysregulated expression of imprinted genes and “2C” genes characteristic of the embryonic totipotent 2-cell state [51, 52]. mESC engineered to lack the TET1-OGT interaction [25] showed a similar global loss of 5mC, apparent in both euchromatic and heterochromatic regions of the genome. Thus, OGT restrains TET activity genome-wide, preventing widespread DNA demethylation and activation of TE expression in heterochromatin.

## RESULTS

### Global decrease in DNA methylation in OGT-deficient mESC

To explore whether OGT affected DNA modification pathways in mESC, we employed an inducible system to disrupt expression of the *Ogt* gene [21]. We cultured mESC (prepared from blastocysts with the genotype *Ogt fl/fl Cre-ERT2 Rosa26-YFP^LSL^* (*Ogt fl*) under non-2i conditions (KOSR serum replacement media with LIF) and treated them for 6 days with 4-hydroxytamoxifen (4-OHT) to generate *Ogt iKO* mESC; *Ogt fl* cells not treated with 4-OHT were used as controls (*Ctrl*) (**Fig. 1A**). Since OGT is essential for cell viability [12, 21], our system of complete *Ogt* gene deletion [21] is superior to OGT depletion with siRNA, shRNA or CRISPR/Cas9 technology, which result in selection for cells that escape depletion and continue to maintain some level of OGT. We sorted cells expressing high levels of YFP (for gating strategy, see **Suppl. Fig. S1A, S1B**) and confirmed, consistent with our previous study [21], that *Ogt iKO* mESC displayed an almost total loss of OGT protein and the *O*-GlcNAc modification (**Fig. 1B**), *Ogt* mRNA by RNA-seq (**Suppl. Fig. S1C**), and OGT as well as OGA by mass spectrometry (**Suppl. Fig. S1D**).

**Figure 1:**
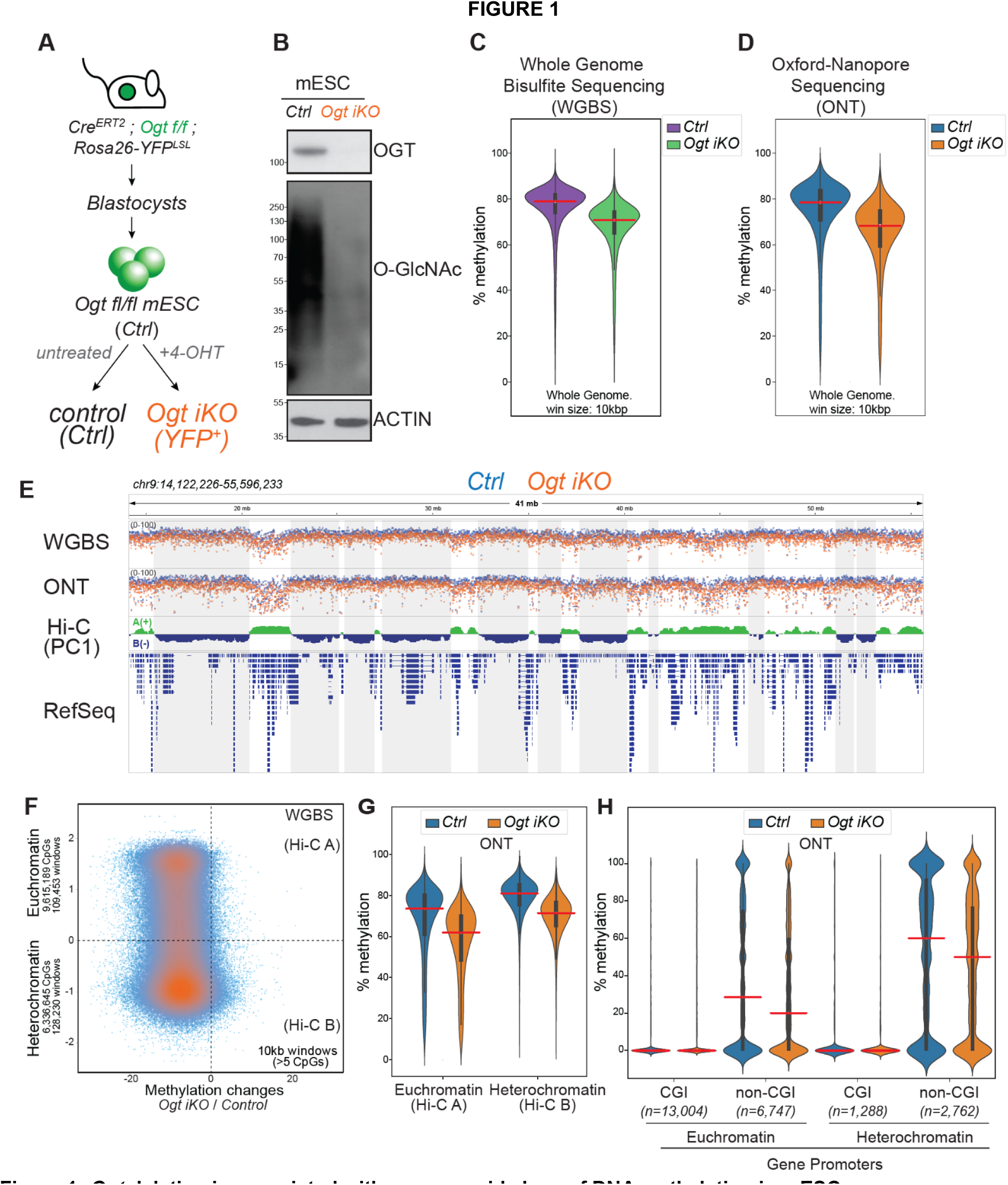
*Ogt* deletion is associated with genome-wide loss of DNA methylation in mESC. **A**, Flow-chart for generation of *Ogt iKO* mESC. *Ogt flfl, Cre^ERT2^, Rosa26-YFP^LSL^* mESC were cultured for 6 days with 4-OHT to yield *Ogt iKO* mESC. Sorted YFP^+^ *Ogt fl* mESC were used for all experiments*. Ogt fl* mESC not exposed to 4-OHT were used as control (*Ctrl*, *Ogt fl*). **B**, Western blots showing loss of OGT protein (*top*) and the *O*-GlcNAc modification (*bottom*) in whole-cell lysates prepared after 6 days of exposure of *Ogt fl* mESC to vehicle or 4-OHT. Actin was used as a loading control. **C-D**, Violin plots showing global loss of DNA methylation in *Ctrl Ogt fl* mESC and *Ogt iKO* mESC measured by Whole-Genome Bisulfite Sequencing (WGBS; **C**) and Oxford Nanopore Technologies sequencing (ONT, **D**). DNA modification (5mC+5hmC) levels were assessed in 10 kb windows across the genome. **E**, Genome browser view of a portion of Chr. 9, showing DNA modification levels (5mC+5hmC) in all CpGs with sufficient coverage in *Ctrl Ogt fl* (*blue*) and *Ogt iKO* (*orange*) mESC. Overlaid tracks of WGBS (*top track*) and ONT (*2^nd^ track*) sequencing data are shown. HiC PC1 values (*3^rd^ track*) were used to identify euchromatin (positive values, Hi-C A compartment, *green*) and heterochromatin (negative values, Hi-C B compartment, *blue*). The majority of CpGs lose DNA modification (5mC+5hmC) in *Ogt iKO* compared to *Ctrl Ogt fl* mESC. **F**, Dot plot of change in DNA methylation (WGBS) in *Ctrl Ogt fl* and *Ogt iKO* in euchromatic (Hi-C A) and heterochromatic (Hi-C B) compartments. Each dot represents a 10 kb window containing at least 5 CpGs covered by at least 5 reads. The majority of CpGs lose DNA modification in *Ogt iKO* compared to *Ctrl Ogt fl* mESC. **G**, Violin plots of DNA methylation (ONT sequencing) in euchromatin (*left*) and heterochromatin (*right*). Mean values are indicated by horizontal red lines. The majority of 10 kb windows lose DNA modification in *Ogt iKO* compared to *Ctrl Ogt fl* mESC. **H**, Violin plots of DNA methylation (ONT sequencing) at proximal gene promoters located in euchromatin (*left*) or heterochromatin (*right*). Promoters of protein coding-genes (−1000 to +500 bp relative to the transcription start site, TSS) were classified as containing or not containing CpG islands (CGI). Mean values averaged over the promoter are indicated by horizontal red lines. The numbers of promoters in each category are indicated. The majority of non-CGI promoters lose DNA modification in *Ogt iKO* compared to *Ctrl Ogt fl* mESC.

Given the connections between OGT and DNA cytosine modification pathways mediated by TETs and DNMTs, we assessed the effects of *Ogt* deletion on DNA methylation in both heterochromatic (Hi-C B) and euchromatic (Hi-C A) genome compartments, defined by calculating Hi-C PC1 values in *WT* mESC [53] (**Fig. 1C-H**, **Suppl. Fig. S1G**). Both whole genome bisulfite sequencing (WGBS) (**Fig. 1C**; **Fig. 1E**, *track 1*; **Fig. 1F**) and Oxford Nanopore Technology long-read sequencing (ONT-seq) [54] (**Fig. 1D**; **Fig. 1E**, *track 2*; **Fig. 1G)** revealed a global reduction of DNA methylation in 10 kb windows across the entire genome in *Ogt iKO* compared to *Ctrl Ogt fl* mESC (**Fig. 1F**, **1G**). We compared DNA methylation levels (ONT-seq) at gene promoters overlapping with CpG islands (CGI promoters), that are associated with housekeeping genes expressed in all cell types, versus non-CGI promoters associated with genes expressed only in specific cell types and located in heterochromatin in non-expressing cell types [55]. DNA methylation at CGI promoters was uniformly low, whereas DNA methylation at non-CGI promoters showed a bimodal distribution of DNA methylation in both euchromatin and heterochromatin and this decreased significantly in *Ogt iKO* compared to *Ctrl Ogt fl* mESC (**Fig. 1H**). DNA methylation decreased on non-CGI gene promoters in *Ogt iKO* versus *Ctrl* mESC (**Fig. 1H**, *compare blue and orange violin plots*).

### Decreased DNMT1 and TET protein levels but increased 5hmC in OGT-deficient mESC

To determine why *Ogt* deletion resulted in a global loss of DNA methylation (5mC+5hmC) by WGBS and ONT, we examined the mRNA and protein levels of DNMT and TET enzymes in *Ogt iKO* mESC by mass spectrometry and western blotting (**Suppl. Fig. S1D-F**). mRNA and protein levels of the maintenance methyltransferase DNMT1 and its partner UHRF1 were both decreased in OGT-deficient compared to control mESC (**Suppl. Figs. S1D, E**), perhaps explaining the loss of DNA methylation over the 6-day time period following 4-OHT exposure. Similarly, mRNA and protein levels of the two major TET enzymes present in mouse ES cells, TET1 and TET2 [56], were also significantly decreased, with an ∼2-fold decrease in TET1 and TET2 peptides in *Ogt iKO* compared to *Ctrl Ogt fl* mESC by mass spectrometry, and a perceptible decrease in TET1 and TET2 protein levels by western blotting (**Suppl. Figs. S1D-F**). There were minor changes, in the protein levels of KAP1 and DNMT3A, and a small increase in DNMT3B levels assessed by western blotting (**Figs. S1D, E**).

Given the tight physical interaction of OGT with TET enzymes and the *O*-GlcNAcylation of TET enzymes by OGT [21–28], we asked if 5hmC levels were altered in *Ogt*-deficient compared to *Ctrl* mESC. Dot blotting of bisulfite-treated DNA, developed with antibodies to cytosine-5-methylenesulfonate (CMS, the product of the reaction of 5hmC with sodium bisulfite [9]) showed a reproducible increase of 5hmC in genomic DNA from *Ogt iKO* compared to *Ctrl Ogt fl* mESC (**Fig. 2A**, **Suppl. Fig. S2A**). Flow cytometry showed that the gain of 5hmC occurred uniformly in the entire population of OGT-deficient mESC, not just in a subpopulation of cells (**Suppl. Fig. S2B**). Notably, the gain of 5hmC occurred despite the substantial decrease in TET1 and TET2 protein levels (**Suppl. Fig. S1D-E**), indicating that OGT maintains TET1 and TET2 protein levels in wildtype mESC but also suppresses TET enzymatic activity. Whole-genome mapping of 5hmC by CMS-IP [57–60] showed that the increase in 5hmC occurred to a similar extent in both euchromatic (Hi-C A) and heterochromatic (Hi-C B) compartments (**Suppl. Fig. S2C, S2D**), even though as expected [60]), 5hmC was predominantly located in euchromatin (regions of high positive or low negative PC1 values) in *Ctrl Ogt fl* mESC. Analysis of CMS-IP data showed that more peaks gained than lost 5hmC (**Suppl. Fig. S2E**).

**Figure 2:**
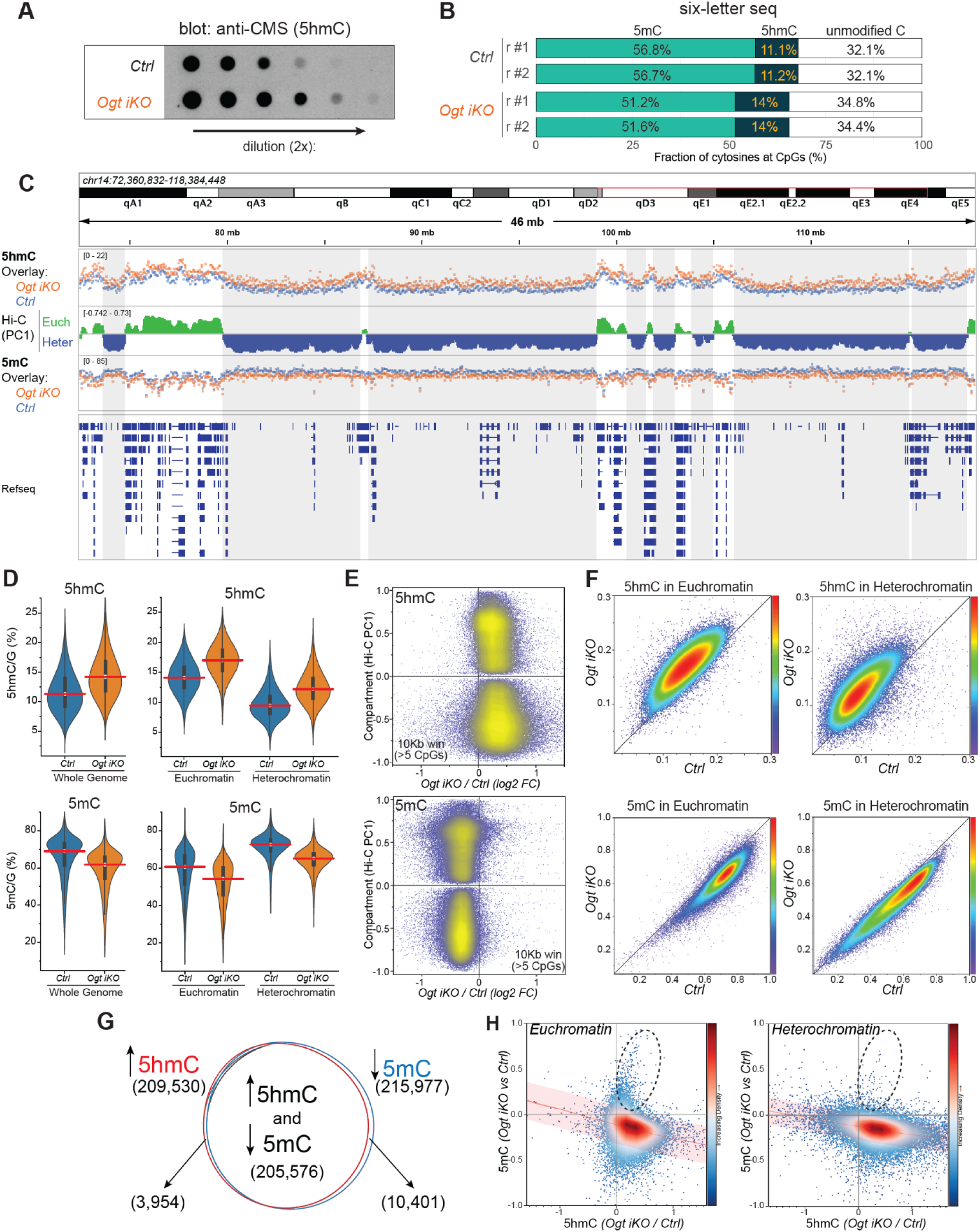
Six-letter sequencing shows that *Ogt* deletion is associated with increased TET activity in mESC. **A**, DNA dot blot showing increased 5hmC in *Ogt iKO* compared to *Ctrl Ogt fl* mESC. A representative experiment is shown; for dot blots of all four replicate experiments please see **Suppl. Fig. 2A**. **B**, Fractions of 5mC, 5hmC and unmodified cytosines in two biological replicates each of *Ctrl Ogt fl* and *Ogt iKO* mESC. **C**, Genome browser view of 5hmC (*top*) and 5mC (*bottom*) in overlaid tracks of *Ogt iKO (orange)* and *Ctrl Ogt fl (blue)* mESC. Values were averaged across 10 kb windows. Hi-C PC1 values (*middle*) distinguish euchromatin (*green*) and heterochromatin (*blue*). *Ogt iKO* (*orange*) mESC exhibit higher 5hmC and lower 5mC in both genomic compartments. **D**, Violin plots showing increased 5hmC (*top*) and decreased 5mC (*bottom*) in *Ogt iKO* compared to *Ogt fl (Ctrl)* mESC at whole genome resolution (*left*) or by chromatin compartments (*right*). **E**, Dot plots showing increased 5hmC (*top*) and decreased 5mC (*bottom*) in *Ctrl Ogt fl* compared to *Ogt iKO* in euchromatin (Hi-C A) and heterochromatin (Hi-C B) compartments. Averaged DNA modification values in 10 kb windows are shown. **F**, Density plots of 5hmC (*top*) and 5mC (*bottom*) in *Ctrl Ogt fl* plotted against *Ogt iKO* mESC in euchromatin (Hi-C A, *left*) and heterochromatin (Hi-C B, *right*) compartments. Averaged DNA modification values in 10 kb windows are shown. The large majority of windows in both compartments show increased 5hmC and decreased 5mC. **G**, Venn diagram showing the overlap between averaged DNA modification values in 10 kb windows that gain 5hmC (blue) and those losing 5mC (red) in *Ogt iKO* compared to *Ctrl Ogt fl* mESC. For the same analysis at the level of individual CpGs, please see **Suppl. Fig. 3E**. **H**, Scatter plots showing the ratio of changes in 5hmC and 5mC in *Ogt iKO* versus *Ctrl Ogt fl* mESC, separated by euchromatin (*top,* Hi-C A) and heterochromatin (*bottom,* Hi-C B) compartments. Averaged DNA modification values in 10 kb windows are shown. The large majority of windows that gain 5hmC in both euchromatin and heterochromatin show loss of 5mC, but a minor subset of windows in euchromatin that gain 5hmC also gain 5mC.

We used six-letter sequencing (seq) [50] to detect both 5hmC and 5mC (**Fig. 2B-H**). Six-letter-seq is an enzymatic single-workflow approach to simultaneously detect both cytosine modifications 5mC and 5hmC, alongside the canonical bases G, C, T and A, at base-resolution in a single sequencing run (for quality control data see **Suppl. Fig. S3A-C**). This method confirmed both the increase in 5hmC and the decrease in 5mC across the genome of *Ogt iKO* compared to *Ctrl Ogt fl* mESC (**Fig. 2B-F**). In *Ctrl* mESC, 5hmC and 5mC are more abundant in euchromatin and heterochromatin respectively (**Fig 2D**, *right panels*, *compare blue (Ctrl) violin plots*); in *Ogt iKO* mESC, there is a clear increase in 5hmC and a clear drop in 5mC in both euchromatic and heterochromatic compartments (**Fig. 2B-F**). The decrease in 5mC largely occurred in the same genomic 10 kb windows as the increase in 5hmC (**Fig. 2G**, **Suppl. Fig. S3D**); the overlap remained substantial (76-80%) when considering individual CpGs (**Suppl. Fig. S3E**). There was an inverse relationship between the gain of 5hmC and loss of 5mC in the majority of 10 kb windows in both compartments (**Fig. 2H**), but a minor subset of regions in euchromatin exhibited concomitant increases of both 5hmC and 5mC in *Ogt*-deficient mESC (**Fig. 2H**, *left panel, dotted oval*). This subset was not observed in heterochromatin, in which the vast majority of regions that gained 5hmC also lost 5mC (**Fig. 2H**, *right panel*).

### The global decrease in DNA methylation in OGT-deficient mESC correlates with increased expression of heterochromatin-localized TEs

*Ogt* deletion resulted in a genome-wide decrease in 5mC (**Figs. 1**, **2, Suppl. Fig. 1-3**), and 5mC suppresses the expression of many TE families [33–39, 54, 61]. We therefore examined the effect of *Ogt* deletion on TE expression (**Fig. 3**). Consistent with their genome-wide loss of 5mC, *Ogt iKO* mESC displayed substantial up-regulation of many TE families, including the youngest L1 families that retain the ability to mobilize (L1 T_F_, G_F_ and A) [62] (**Fig. 3A**). We also observed a striking upregulation of ERVs, notably MERVL-int and MT2_Mm, the internal and regulatory regions of the mouse ERV element ERV-L (MERVL) family, respectively, as well as IAPEy and IAPEz retrotransposons belonging to the IAP family, whose expression is normally repressed at least partially by DNA methylation [33–38] (**Fig. 3A**). By integrating TE expression with data from ONT-seq [54], we observed an overall decrease in DNA methylation at the LTR and 5′UTR sequences of ERVs and L1 families respectively in *Ogt*-deleted mESC (**Fig. 3B**, *violin plots*].

**Figure 3:**
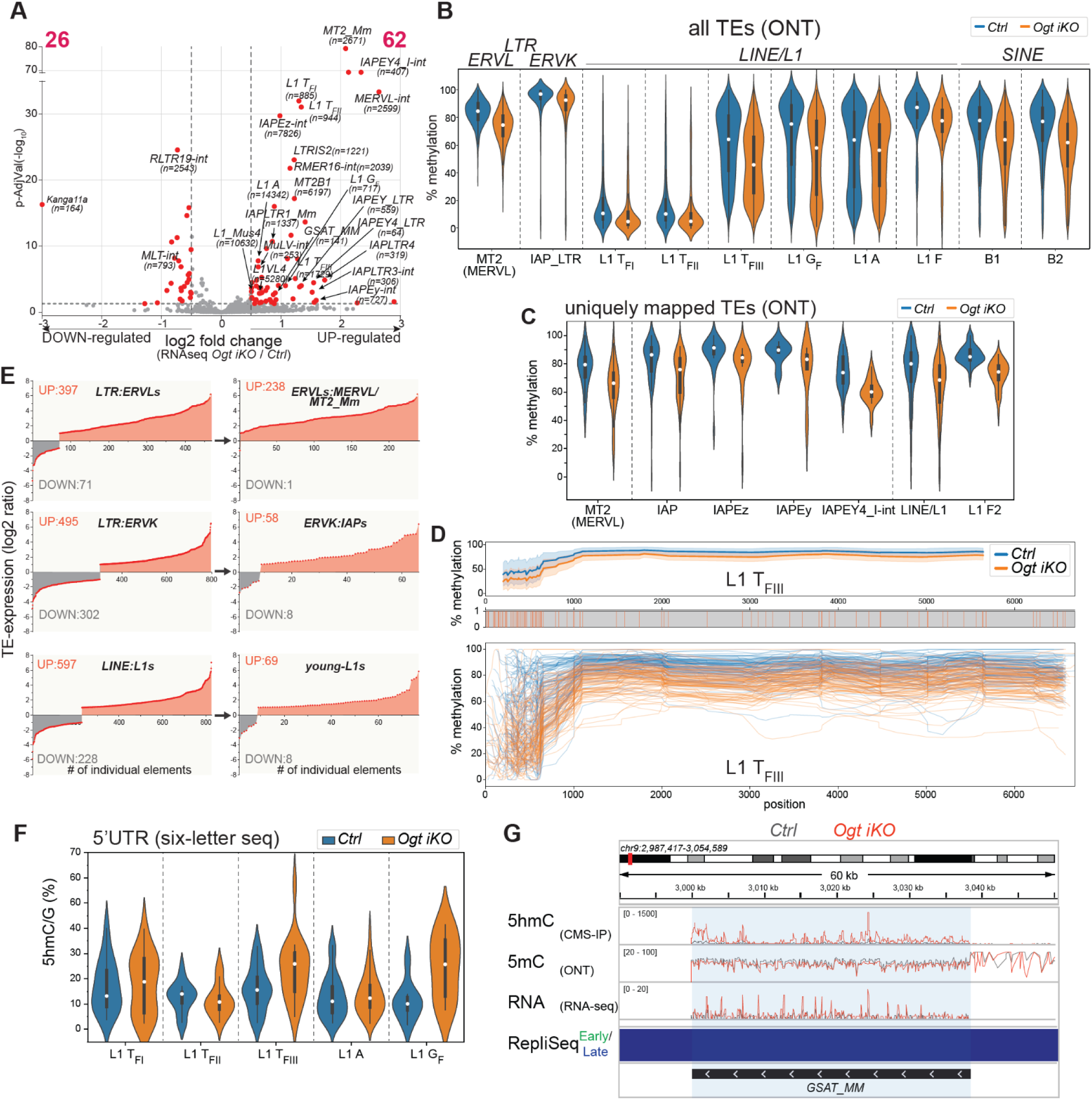
*Ogt* deletion is associated with enhanced expression of LINE-1 and LTR elements in mESC. **A**, Volcano plot of differentially expressed transposable elements (TEs) in *Ogt iKO* relative to *Ctrl Ogt fl* mESC. RNA-Seq data were analyzed for TE expression using the TEtranscripts package to measure the bulk expression of TE subfamilies. Each point represents a specific TE-subfamily; the number of individual copies analyzed for each subfamily is indicated in parentheses. Red dots represent transcripts exhibiting at least a 2-fold change in expression (log2 fold change (log2FC) of +/-1) and a pAdj value <0.05 (calculated by adjusting the p-value with the false discovery rate [FDR]). **B**, CpG methylation levels (5mC+5hmC) ascertained by ONT sequencing of TE subfamilies in *Ogt fl* (blue) and *Ogt iKO* (orange) mESC. Results are shown for MT2 (MERVL promoter), IAP LTRs, the 5′ UTR regulatory regions of LINE-1 elements (L1 A-type, G_F_ and T_F_ of length > 6 kb), and SINE B1 and B2 families. Violin plots indicate the median, interquartile range, and 1.5x interquartile range. **C**, Violin plots showing changes in DNA modification levels measured by ONT sequencing at the regulatory regions of uniquely mapped, differentially expressed TEs in *Ogt iKO* compared to *Ctrl Ogt fl* mESC. **D**, Composite methylation profile of the young L1 T_FIII_ in *Ogt iKO* and *Ctrl Ogt fl* mESC. *Top panel*, *Ogt iKO* mESC (*orange*) show decreased average methylation (5mC+5hmC) along the length of L1 T_FIII_ elements compared to *Ctrl Ogt fl* mESC (*blue*), with particularly pronounced loss of methylation in the 5’ UTR regulatory region at left. *Middle panel*, Red vertical lines indicate the position of CpGs. *Bottom panel*, Methylation profiles of ONT long reads fully covering uniquely mapped L1 T_FIII_ elements. CpGs not confidently called, i.e. absolute (log-likelihood ratio) > 2.5, were omitted. **E**, Analysis of individual uniquely mapped TEs showing dysregulated expression in *Ogt iKO* compared to *Ctrl Ogt fl* mESC. RNA-seq data were analyzed to determine the expression of individual differentially expressed TEs using TE-transcripts and DESeq2 (log2 fold change (log2FC) of +/-1 and p-value <0.05). The number of up- (*red*) or down-regulated (*gray*) unique TEs are indicated in each case. The majority of differentially expressed and uniquely mapped TEs in the ERVL LTR (397/468, 85%), ERVK LTR (495/797, 62%) and L1 (597/825, 72%) subfamilies (*left panels*) were upregulated in *Ogt iKO* compared to *Ctrl Ogt fl* mESC, and this tendency for increased expression was even more pronounced in the youngest members of each family (*right panels*): MERVL-int/MT2 subfamily (238/239, 99%); IAP subfamilies (58/66, 88%); young LINE-1 elements (L1 T_FI, II, III_, L1 A, L1 G_F_) (69/77, 90%). **F**, Violin plots showing changes in 5hmC levels, measured by six-letter-seq, at the indicated 5’UTR of the L1 subfamilies in *Ogt iKO* mESC compared to *Ctrl Ogt fl* mESC. **G**, Genome browser view showing increase in 5hmC (CMS-IP) and reduction of 5mC (ONT seq) at GSAT_MM major satellite regions, concomitantly with increased numbers of RNA-seq reads mapping to these repetitive regions. The increased expression of GSAT_MM in *Ogt iKO* mESC was confirmed using total ribo-depleted RNA-seq (*not shown*).

Conventional analysis of TE expression involves aligning RNA-seq reads while permitting multi-mappers over hundreds or thousands of individual TEs that are very similar in sequence [63], allowing the mean expression of each TE subfamily to be estimated (**Fig. 3A**). However, this method does not provide specific information about the expression of individual TE loci in the family. To relate TE expression to 5mC, 5hmC and other genomic features, and to ask whether increased TE expression was linked to increased expression of nearby genes, we identified the subgroup of individual TEs within each subfamily that showed increased expression in *Ogt iKO* compared to *Ogt fl* mESC (**Fig. 3C**, **Fig. 3E**; also see *Methods* and **Fig. 4**). The loss of DNA methylation in *Ogt iKO* versus *Ctrl Ogt fl* mESC was accentuated in this smaller subset of uniquely mapped TEs (**Fig. 3C**). As an example, the 5′UTR regulatory regions of the L1 T_FIII_ subfamily showed an average reduction in DNA methylation compared to the youngest mobile L1 T_FI/II_ subfamilies [62] whose 5’ UTRs are very poorly methylated (**Fig. 3B**; **Fig. 3D**, *upper panel*). In contrast, individual uniquely mapped TEs in the L1 T_FIII_ subfamily showed a wide variation in DNA modification, especially at their 5’UTRs, in both *Ctrl* and *Ogt iKO* mESC (**Fig. 3D**, *lower panel*). Moreover, in each upregulated TE family, only a small number of uniquely mapped TEs showed decreased expression compared to the larger number of TEs showing increased expression in *Ogt iKO* versus control *Ogt fl* mESC (**Fig. 3E**).

**Figure 4:**
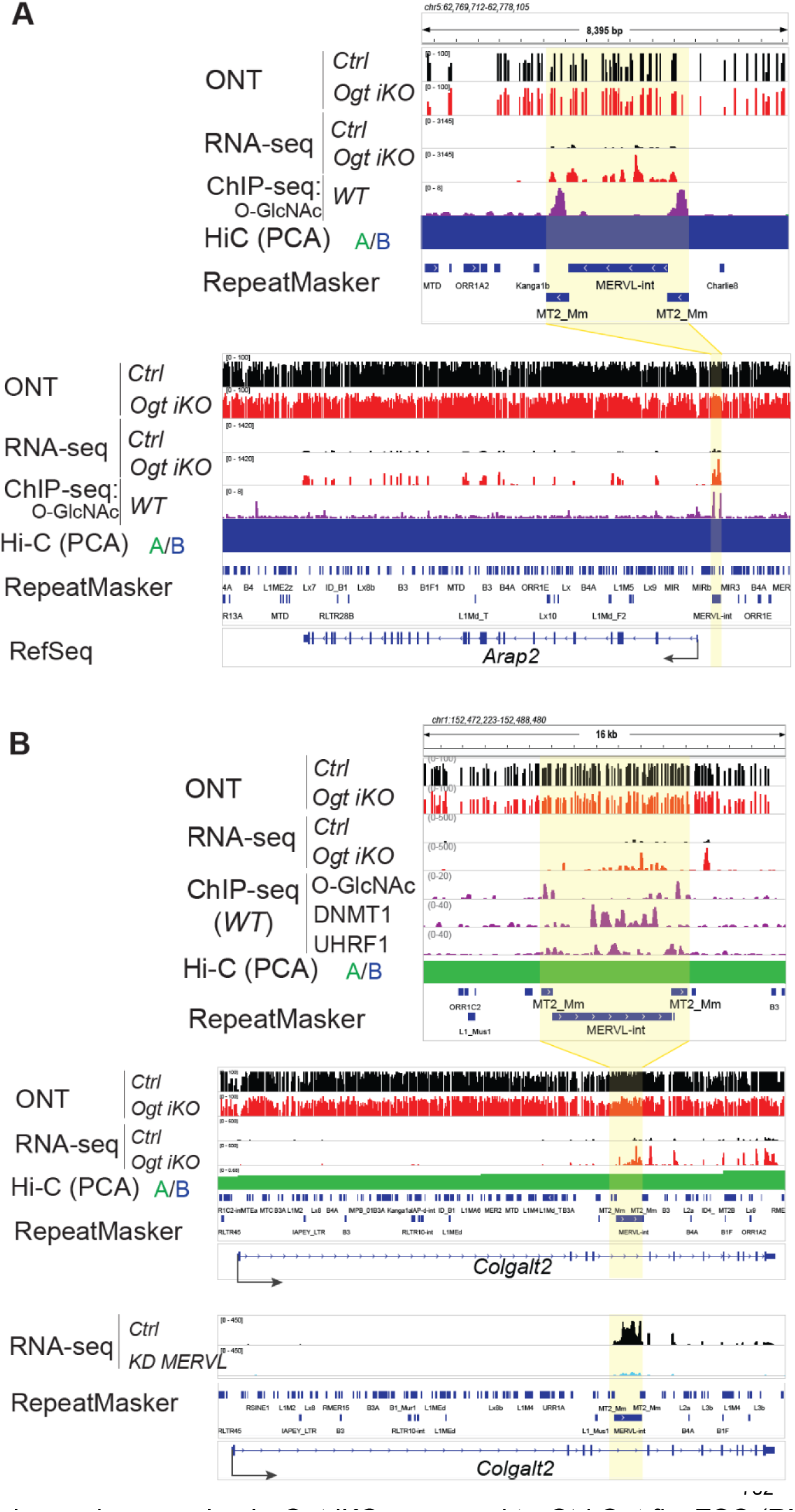
Increased expression of MERVL-int/MT2_Mm TEs is associated with increased expression of 3’ genes and exons in *Ogt iKO* mESC. In all figures, the top four tracks show ONT-seq and RNA-seq data from *Ctrl* and *Ogt iKO* mESC, followed by ChIP-Seq for *O*-GlcNAc, DNMT1 or UHRF1 as indicated in WT mESC. *Bottom track*, HiC data (PC1) indicate the presence of these regions at euchromatin (green) or heterochromatin (blue). **A,** *top*. Genome browser view of a uniquely mapped MERVL-int/MT2 element 5’ of the *Arap2* gene in heterochromatin. Note the loss of DNA modification (ONT sequencing, 5mC+5hmC) at certain CpGs (*ONT, top tracks*), the increased expression of the MERVL elements in *Ogt iKO* compared to *Ctrl Ogt fl* mESC (RNA-seq tracks), and the presence of the *O*-GlcNAc modification on chromatin-associated proteins occupying the two MT2_Mm regulatory regions, based on *O*-GlcNAc ChIP-seq in mESC [33]. *bottom*, Activation of the MERVL-int/MT2 element shown in A correlate with enhanced expression of *Arap2*. The MERVL TE and the *Arap2* gene are encoded on the same strand, suggesting *cis* regulation of *Arap2* by the MERVL TE, acting either as a promoter or as a nearby enhancer. **B,** *top*. A MERVL-int/MT2_Mm TE located in an intronic region of the *Colgalt2* gene located in euchromatin is up-regulated in *Ogt iKO* compared to *Ctrl Ogt fl* mESC (RNA-seq tracks). Note the loss of DNA modification (ONT-seq) at most CpGs (*ONT, top tracks*); the presence of the *O*-GlcNAc modification on chromatin-associated proteins occupying the two MT2_Mm regulatory regions, based on *O*-GlcNAc ChIP-seq data in mESC [33]; and occupancy of the element by Dnmt1 and Uhrf1, based on ChIP-seq data in mESC [66]. *Middle*, Only the exons of *Colgalt2* located 3’ of this upregulated MERVL-int/MT2_Mm display enhanced expression in *Ogt iKO* compared to *Ctrl Ogt fl* mESC (RNA-seq tracks). The MERVL TE and the *Colgalt2* gene are encoded on the same strand, suggesting *cis* regulation of the 3’ exons of the *Colgalt2* gene by the MERVL TE, acting either as a promoter or as a nearby enhancer. *Bottom*, Knockdown (KD) of MERVL-int elements causes reduced expression of the MERVL-int TE contained in the intron of the *Colgalt2* gene, as well as reduced expression of the *Colgalt2* gene itself. Data from early embryos with MERVL-int KD as well as its control counterparts [73] were reanalyzed.

5hmC is most highly enriched in gene bodies of the most highly transcribed genes and at the most active enhancers [57–60]. To ask if this was also true for expressed, uniquely mapped TEs, we used data from six-letter-seq to examine the loss of 5mC and gain of 5hmC respectively at the regulatory regions (**Fig. 3F**) and gene bodies (**Suppl. Fig. 3F**) of these TEs. We confirmed the loss of 5mC (**Fig. 2D-F**) and observed a gain of 5hmC (**Fig. 3F**) in the 5’ UTRs of nearly all young L1 subfamilies (defined as longer than 6 kb and labelled young-L1 in **Fig. 3E**) in *Ogt iKO* relative to *Ctrl Ogt fl* mESC. The gain of 5hmC in L1 5ʹUTRs was mainly driven by gains in the T_FI_, T_FIII_ and G_F_ subfamilies after *Ogt* deletion (**Fig. 3F**). The family of GSAT_MM satellite elements also showed increased 5hmC and decreased 5mC (**Fig. 3G**) concomitantly with increased expression (**Fig. 3A**). Together the data indicate that OGT is required to suppress expression of many L1 and LTR elements by preventing the increase of 5hmC in the transcribed units and regulatory regions of TEs in normal mESC.

We confirmed increased TE expression by qRT-PCR (**Suppl Fig. S4A**) as well as mass spectrometry (**Suppl. Fig. S4B**). *Ogt iKO* cells contained increased levels of peptides derived from LINE-1 ORF-1p and ORF-2p proteins, and from GAG proteins encoded in MERVL-int and IAP elements in *Ogt iKO* versus *Ogt fl (Ctrl)* mESC (**Suppl. Fig. S4B**). Moreover, expression of a MERVL reporter [51] stably integrated into *Ogt fl* mESC was uniformly increased in the *Ogt iKO* mESC population after *Ogt* deletion (**Suppl. Fig. 4C**); the rightward shift of the entire population represents an increase in baseline MERVL expression in all *Ogt iKO* cells, rather than an increase in the small population of MERVL reporter-positive cells (not apparent in this experiment) that cycle in and out of the 2-cell (2C) state in wildtype mESC [51]. Finally, the *O*-GlcNAc modification was enriched at the regulatory regions of mouse MERVL, L1, and ERVK subfamilies of transposable elements in wildtype mESC (**Suppl. Fig. 4D**). Thus, increased expression of these TEs after *Ogt* deletion might reflect loss of 5mC and/or loss of the *O*-GlcNAc modification in chromatin proteins binding TE regulatory regions [33].

The TE families that were most strikingly upregulated in *Ogt iKO* relative to *Ogt fl* mESC – MERVL, IAPs and young L1 T_F_, G_F_ and A elements – corresponded to those that were predominantly located in heterochromatic regions of mESC (**Suppl. Fig. 4E**). TEs whose regulatory regions were enriched for *O*-GlcNAc (**Suppl. Fig. 4D**) were even more enriched in heterochromatin compared to all TEs (**Suppl. Fig. 4F**).

We used ONT-seq to investigate an IAPEy element (annotated with two LTR flanking an adjacent internal (int) region) to a unique location in the euchromatic region of the genome (**Suppl. Fig. S5A**, *top two tracks*). This IAPEy element was poorly expressed in *Ctrl Ogt fl* mESC but highly expressed in *Ogt iKO* mESC (**Suppl. Fig. S5A**, *RNA-seq tracks*), and displayed clear evidence of DNA demethylation at both ends of the element, especially the 5’ LTR (**Supp. Fig. S5A**, *yellow shaded rectangles*). Reanalysis of published data [33, 64–67] showed that this region was occupied by chromatin proteins bearing the *O*-GlcNAc modification, as well as by OGT, DNMT1 and UHRF1 (**Supp. Fig. S5A**, *ChIP-seq tracks*). The data suggest that OGT suppresses the expression of this IAP element by modifying chromatin-bound transcriptional regulators at its LTRs, and recall a previous finding in which targeting OGA as a deadCas9 fusion to the promoter regions of another subfamily of IAP elements (IAPEz) to remove the *O*-GlcNAc modification resulted in a dramatic and selective reactivation of IAPEz expression [33]. Similarly, an MT2_Mm regulatory region and a relatively young L1Md_F3 element located adjacent to one another in heterochromatin were suppressed in *Ctrl Ogt fl* mESC but expressed in *Ogt iKO* mESC; the 5’ UTR of the L1Md_F3 element corresponded to a peak of *O*-GlcNAc modification and displayed demethylation at certain CpGs (**Supp. Fig. S5B**, *yellow shaded rectangles*).

### Increased TE expression in OGT-deficient mESC correlates with increased expression of genes and exons located 3’ of the expressed TE

TEs are known to influence the expression of nearby genes, in part because their regulatory elements can act as enhancers and promoters in *cis* [36–38, 68–72]. We asked whether TE expression was associated with altered expression of nearby genes (**Fig. 4**). Indeed, expression of the *Arap2* gene and a MERVL-int/MT2_Mm element located 5’ of *Arap2* were both upregulated in *Ogt iKO* compared to *Ctrl* mESC, concomitantly with altered DNA methylation of certain CpGs in the MT2_Mm regulatory (LTR) region just 5’ of the annotated MERVL-int TE (**Fig. 4A**). In another example, increased expression of a MERVL element (MERVL-int with two flanking MT2_Mm LTRs) located in an intron of the *Colgalt2* gene (**Fig. 4B**, *top*) was linked to increased expression of only those coding exons of *Colgalt2* that were located 3’ of the MERVL element, with no change in expression of exons located 5’ of the element (**Fig. 4B**, *middle*), consistent with *cis*-regulation of the 3’ but not 5’ exons of *Colgalt2* by the expressed MERVL TE. This finding was confirmed by analyzing data from a separate study [73], showing that early mouse embryos express this MERVL element as well as the 3’ (but not the 5’) exons of *Colgalt2*, and that global MERVL knockdown (KD) in early embryos resulted in reduced expression of both this particular MERVL element and the 3’ exons of *Colgalt2* (**Fig. 4B**, *bottom*). In both examples, the MERVL regulatory regions were marked with *O*-GlcNAc, potentially linking loss of the modification to increased TE and gene/exon expression in *Ogt iKO* mESC (**Fig. 4**). Thus, expression of MERVL elements is normally suppressed, directly or indirectly, by OGT in early embryos, and the activation of these elements in early embryos and in OGT-deficient mESC is associated with increased expression of the *Arap2* gene and the exons of *Colgalt2* located 3’ of the MERVL element. The MERVL elements may behave as alternative promoters, forming fusion transcripts with genes and exons that are located 3’ of the element [68], or may function as proximal enhancers to activate 3’ genes and exons in *cis*.

*Ogt iKO* mESC showed upregulation of many genes (1661 up-regulated and 749 down-regulated genes), including a large number (43/52) of 2C-like genes [51, 52] compared to *Ogt fl* (*Ctrl*) mESC (**Suppl. Fig. S6A**). The majority of imprinted genes (73/90) were also upregulated, but most genes encoding the rapidly evolving KRAB ZFPs [43, 44] (64/88) were downregulated in *Ogt iKO* compared to *Ogt fl/fl* mESC (**Suppl. Fig. S6B, C**). OGT deletion was also associated with a striking upregulation of Interferon-Stimulated Genes (ISGs) (**Suppl. Fig. S6D**). Gene Set Enrichment Analysis (GSEA) revealed enrichment for Inflammatory Response pathways and Response to Viruses in *Ogt iKO* compared to *Ctrl Ogt fl/fl* mESC (**Suppl. Fig. S6E-F**). These data suggest a potential involvement of OGT in nucleic acid sensing and antiviral response pathways, and recall the connections among heterochromatic DNA hypomethylation and increased TE expression in cancer, inflammation, cellular senescence and ageing [39, 74–77].

### Blocking the TET-OGT interaction mimics the global loss of DNA methylation observed in OGT deficient cells

The D2018 residue of TET1 is essential for the TET1-OGT interaction, and mESC bearing a single point mutation, D2018A, in the *Tet1* gene showed markedly decreased DNA methylation levels (5mC+5hmC, by WGBS) [25]. This decrease was observed in both euchromatic and heterochromatic compartments (**Fig. 5A-C**), with the greatest loss of DNA methylation observed in the most highly heterochromatinized 10 kb windows (most negative Hi-C PC1 values, **Fig. 5C**). The decrease of 5mC+5hmC was associated with the same type of striking loss of 5mC at TEs (**Fig. 5D**) as shown here by ONT and six-letter sequencing for OGT-deficient mESC (**Figs. 1**, **2**). Thus, blocking the TET1-OGT interaction with a single point mutation has the same effect of reducing global DNA methylation as OGT deficiency itself.

**Figure 5:**
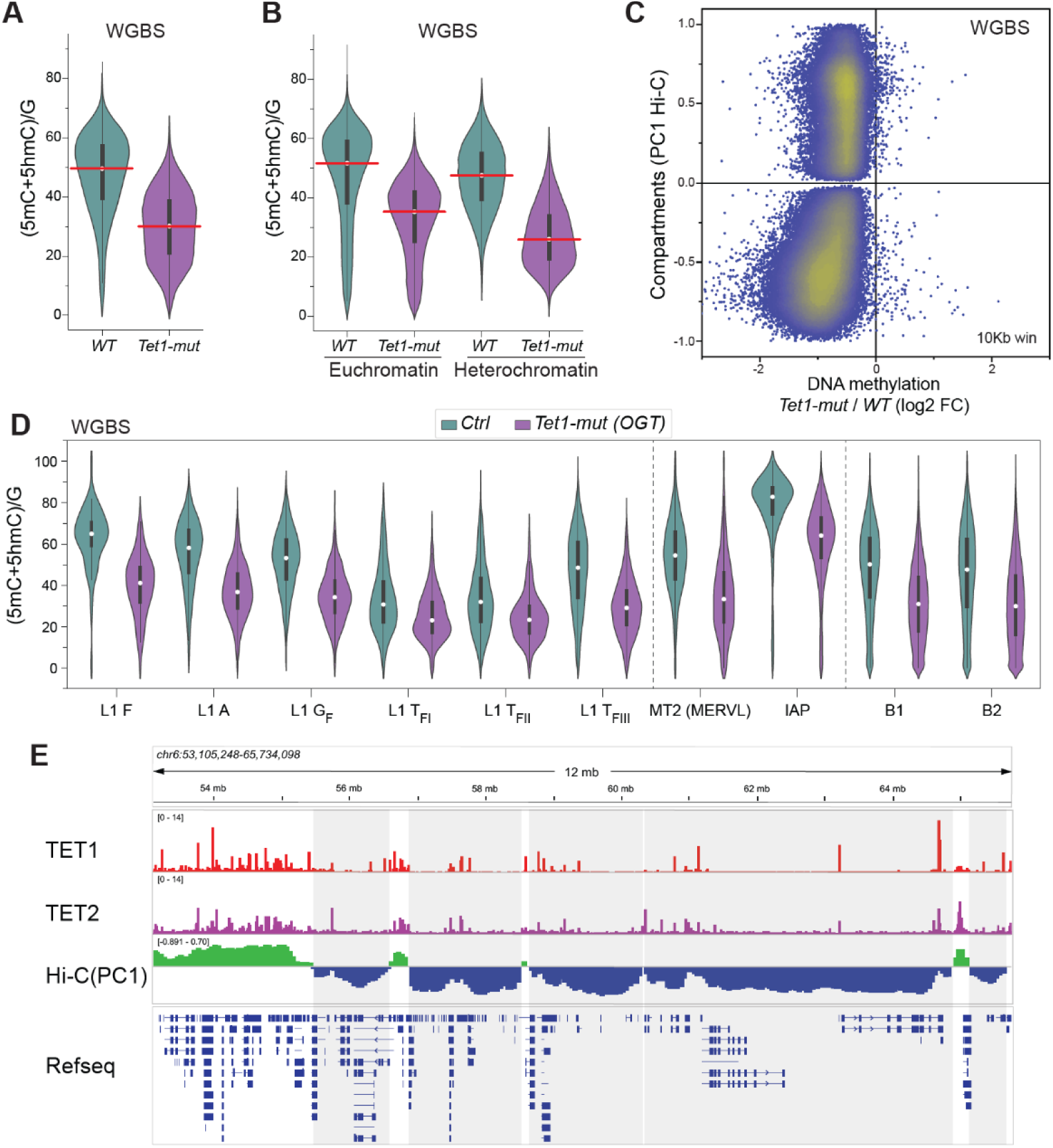
Disruption of the TET1-OGT interaction induces a genome-wide loss of DNA methylation in mESC. Reanalysis of WGBS data from mESC bearing a single point mutation (D2108A) in Tet1 that disrupts the TET1-OGT interaction [25]. **A**, Violin plots of DNA methylation (WGBS, 5mC+5hmC) in the whole genome using 10 kb windows. **B**, Violin plots of DNA methylation (WGBS, 5mC+5hmC) levels in euchromatin (*left*) and heterochromatin (*right*) in *WT* and Tet1 D2108A *(Tet1-mut*) mESC. Mean values are highlighted with a white dot. **C**, Dot plot of changes in DNA methylation (WGBS, 5mC+5hmC) in 10 kb windows in *WT* and *Tet1-mut* mESC in HiC-A and B compartments. **D**, DNA methylation (5mC+5hmC) levels at indicated TE subfamilies in *WT* and *Tet1/OGT-mut* mESC. **E**, TET1 and TET2 proteins bind largely euchromatic regions (positive or low negative PC1 values) in WT mESC*Top two tracks*, Genome browser view of reanalyzed ChIP-seq data for TET1 and TET2 [64, 65] from WT mESC. *Bottom track*, Hi-C PC1 data to identify euchromatin and heterochromatin compartments.

We evaluated the genomic distribution of TET1 and TET2 in heterochromatin of wildtype mESC (**Fig. 5E**, **Suppl Fig S7**) by reanalyzing prior ChIP-seq data [64, 65]. The analysis confirmed that TET1 and TET2 are both located primarily in euchromatin (Hi-C A compartment with positive PC1 values) in mESC, or at regions of relatively low negative PC1 values that – with higher resolution Hi-C or micro-C mapping – might be shown to be located in euchromatin (**Fig. 5E**). This distribution is consistent with our finding that the bulk of 5hmC in mESC and other cell types is located in euchromatin, at the gene bodies of highly expressed genes and at active enhancers [57–59]. A relatively small fraction (12%) of both TET1 and TET2 occupied heterochromatic regions (Hi-C B compartment, with the caveat that the resolution of our Hi-C was ∼50 kb) (**Suppl. Fig. S7A**); however, because TET1 had about twice as many ChiP-seq peaks overall as TET2 (9582 versus 4413; **Suppl. Fig. S7A**), consistent with its higher expression in mESC [21], the number of TET1 ChIP-seq peaks in heterochromatin was also twice that of TET2 (**Suppl. Fig. S7B**). These data, and our prior reanalysis [60] of data from [78], suggest that TET1 may have a more prominent role than TET2 in maintaining high 5hmC and 5mC levels at heterochromatic regions of the genome.

### Identification of proteins that interact with endogenous OGT

There have been numerous analyses of OGT-interacting proteins by mass spectrometry, but the assays have primarily been performed in various transformed cell lines or overexpression of exogenous versions of OGT [28]. To focus on proteins that interact robustly with endogenous OGT expressed at physiological levels in mESC, we used homology-directed repair to generate two mESC cell lines in which a 3x-FLAG or a StrepTag epitope tag had been “knocked-in” by homology-directed repair into the first coding exon of the endogenous *Ogt* gene (**Fig. 6A, B**). Quantitative mass spectrometry of nuclear proteins that co-immunoprecipitated under native conditions with the anti-FLAG antibody from the FLAG-OGT but not the StrepTag-OGT mESC line identified many known OGT interactors, including several known components of epigenetic regulatory complexes, including the NuRD complex which contains MBD3, HDAC1 and HDAC2 and is known to interact with TET1 [22, 80]; HCF1 (encoded by HCFC1), a component of MLL/COMPASS complexes [12]; PROSER1, an adapter protein that modulates TET2 *O*-GlcNAcylation and regulates the methylation status of a small subset of CpG-rich enhancer regions [29]; and the ASXL1-BAP1 polycomb response-deubiquitinase (PR-DUB) complex [81]. OGT also co-immunoprecipitated with many repressive heterochromatin-associated proteins including KAP1; the KRAB-ZFP proteins ZFP57 and ZFP655; and subunits of the PRC2 complex (SUZ12, EED, JARID2) (**Fig. 6C**). We also detected a few interactions not reported previously, for instance with DNMT1 and with USP7, a deubiquitinase that controls p53/MDM2 stability [82, 83] and has also been implicated in regulating the stability and function of the DNMT1/UHRF1 complex [84, 85] (**Fig. 6C-D**). Of these, only DNMT1, TET1, TET2 and PROSER1 were constitutive substrates of OGT, judged by *O*-GlcNAc modification [38, 39, 41–45] (**Fig. 6E** and *not shown*).

**Figure 6:**
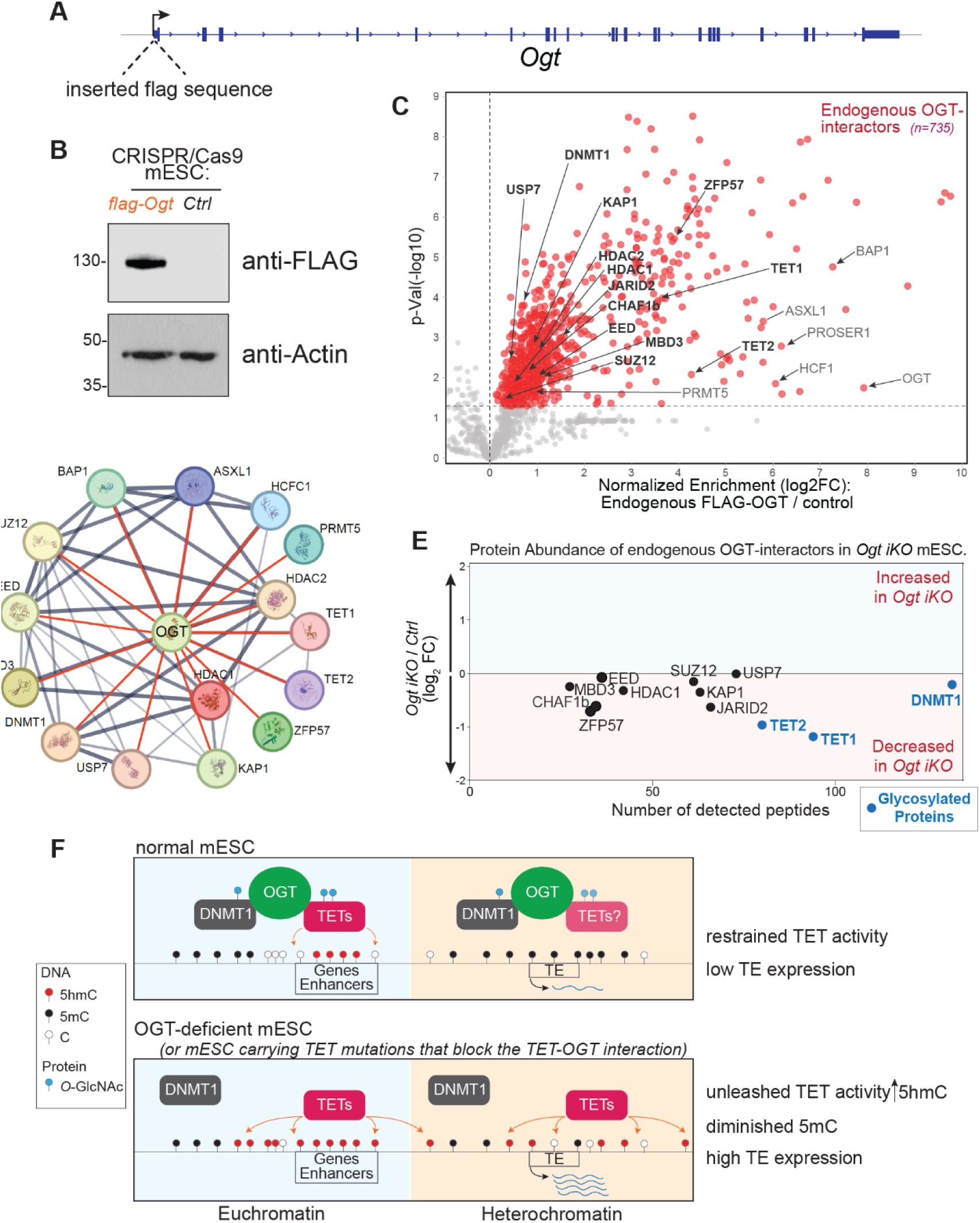
OGT expressed at endogenous levels interacts with many heterochromatin-associated proteins in mESC. **A**, Strategy to generate flag-tagged *Ogt* mESC. A sequence encoding the FLAG epitope tag was knocked-in into the first exon of the *Ogt* gene in wild type mESC using CRISPR/Cas9-assisted homology-directed repair, resulting in the presence of a FLAG tag at the N-terminus of endogenous OGT protein. **B**, Western blot analysis of *FLAG-Ogt* and wildtype mESC. Expression of FLAG-OGT protein was detected using anti-FLAG-antibodies. ACTIN was used as loading control. **C**, Volcano plot highlighting heterochromatin-associated proteins, including DNMT1, TRIM28, ZFP57, USP7 and HDAC1 and 2, that co-immunoprecipitated with FLAG-OGT from nuclear lysates and were identified by mass spectrometry. mESC expressing a StrepII tag and immunoprecipitated with the same anti-FLAG antibody served as a negative control. Many known OGT interactors, including HCFC1, TET1 and TET2, ASXL1 and BAP1 were also detected in our analyses. **D**, STRING diagram showing coimmunoprecipitation of chromatin-associated proteins with OGT. Known interactions identified in human cancer cell lines are shown in gray; interactions identified in mESC in this study are shown in red. **E**, TET1, TET2 and DNMT1 are *O*-GlcNAcylated (blue dots) and show decreased expression in *Ogt iKO* compared to *Ctrl Ogt fl* mESC. **F**. Model illustrating the increase of TET activity in both euchromatin and heterochromatin after *Ogt* deletion. *Ogt iKO* mESC lose *O*-GlcNAcylation and show a substantial decrease of DNMT1 and UHRF1. The genome-wide constraint imposed by OGT on TET activity in WT mESC is unleashed upon OGT deletion, leading to a rapid genome-wide increase in 5hmC and concomitant decrease of 5mC in both euchromatic and heterochromatic compartments. The decrease of 5mC at the regulatory and transcribed regions of TEs, particularly those in heterochromatin, is accompanied by a striking increase in TE expression in OGT-deficient mESC.

Reanalysis of published ChIP-seq data [22, 33, 64–67, 86] showed that OGT, the *O*-GlcNAc modification and the TET/DNMT proteins TET1, TET2, DNMT1 and DNMT3A co-occupied the 5’ UTR regions of the young silenced LINE1 subfamilies L1 TF_III_ and L1_A_ in WT mESC (**Suppl. Fig. S8**), consistent with the possibility that these proteins complex with OGT at TE regulatory regions to control or modulate TE expression. Together, our data confirm a broad role for OGT in repressing TE expression and sustaining the normal properties and function of heterochromatin.

## DISCUSSION

We show here that OGT maintains DNA methylation genome-wide, in a manner dependent on its interaction with TET1 (**Fig. 6F**). We used a system that allowed for complete deletion of the Ogt gene. Acute deletion of the *Ogt* gene in mESC resulted in a global increase in 5hmC and a global reduction of 5mC in both euchromatic and heterochromatic compartments, exactly as expected for increased TET enzymatic activity. Notably, mESC engineered to have a single point mutation in TET1 that abrogated the TET1-OGT interaction showed a global loss of DNA methylation, very similar to that observed in OGT-deficient mESC [25]. Thus, OGT globally restrains the enzymatic activity of TET proteins, extending previous reports that all three TETs bind tightly to OGT [22–24] and that TET1 recruits OGT and TET2 to chromatin [22]. Our studies emphasize the striking connections among OGT, DNA methylation/demethylation pathways mediated by TET1 and DNMT1/UHRF1, and TE suppression mediated by KAP1 in mESC.

The functional crosstalk among TET, OGT and DNMT1 enzymes manifests in diverse ways. OGT suppresses TET activity in mESC (this study), but TET proteins may potentiate OGT activity based on a reported ∼40% decrease in global *O*-GlcNAc levels in Tet2-deficient compared to control bone marrow [24]. OGT requires DNMT1 to bind KAP1 and maintain TE suppression [33], but *Ogt* deletion resulted in decreased levels of DNMT1/UHRF1 (this study), and DNMT1 catalytic activity was reduced in HepG2 cells cultured in high glucose and the OGA inhibitor Thiamet-G [87]. It will be important in future investigations to determine if different *O*-GlcNAc modifications on TET1, TET2 and DNMT1 make distinct contributions to protein stability and function.

As expected [88], the consequences of losing DNA methylation after *Ogt* deletion were apparent primarily in heterochromatin (**Fig. 6F**). Decreased heterochromatic DNA methylation in *Ogt*-deficient mESC compromised heterochromatin integrity, and resulted in rapid TE de-repression within 6 days of exposure to 4-OHT. Increased 5hmC and decreased 5mC were observed in all classes of transposable elements (LINEs, SINEs and LTRs), but increased TE expression after OGT deletion was most striking in young L1 T_F_ and LTR (MERVL, IAP /ERVK) elements that are predominantly located in heterochromatin, and whose regulatory regions are bound by *O*-GlcNAcylated proteins in wildtype mESC. We note, however, that expression of certain L1 subfamilies may depend less on DNA methylation than on H3K9me2/3 methylation [86]. Our data corroborate and extend a previous study showing that partial OGT depletion with shRNA in mESC resulted in increased TE expression, an effect partly reversed by simultaneous depletion of TET1 [45]. In contrast, most SINEs (e.g. B1, B2), which are preferentially contained in euchromatin, showed increased 5hmC but were poorly expressed in *Ogt iKO* mESC. SINEs are often present in transcribed units (e.g. introns of protein-coding genes); hence the significant increase of 5hmC in SINEs in the absence of increased gene expression could reflect the well-known gain of 5hmC at the gene bodies of highly expressed genes [58, 59]. The data emphasize the complexity of TE regulation by DNA and H3K9 methylation/demethylation pathways in mESC.

Analysis of the gene neighborhood of individual uniquely expressed TEs showed that in certain cases, increased TE expression in OGT-deficient mESC correlated with increased *cis*-expression of 3’ genes and exons [66]. TEs can provide alternate transcription start sites [68], and TE regulatory regions can act as enhancers in *cis* [69, 89]. Either mechanism could underlie the selective increase in transcription of the *Arap2* gene and the exons 3’ of the MERVL element in the *Colgalt2* gene. TE transcripts can also be spliced ectopically into existing transcripts [36, 72], and there is a strong connection of acute OGT inhibition with changes in the levels and *O*-GlcNAcylation of splicing factors as well as “detained intron” retention [90]. Overall, it is clear from our own and many other studies that even in the absence of retrotransposition, increased TE expression is likely to be associated with diverse, stochastic alterations in gene expression patterns in cells.

An unexpected consequence of OGT deficiency was increased ISG expression and enrichment of anti-viral response pathways in *Ogt iKO* compared to *Ctrl Ogt fl/fl* mESC. The underlying mechanism is not clear, but likely involves an innate immune response to heterochromatic DNA hypomethylation and increased TE expression. Treatment of cancer cells with the DNMT inhibitors 5-azacytidine and decitabine – which also results in global reduction of DNA methylation – correlates with increased expression of transposable elements, increased levels of cytoplasmic nucleic acids and increased ISG expression (“viral mimicry”) [91, 92]. The data suggest a prominent role for the TET-OGT interaction in suppressing “sterile” inflammation (i.e. inflammation in the absence of pathogen infection) by maintaining physiological levels of DNA methylation and normal patterns of gene and TE expression across the mESC genome.

OGT-deficient cells showed a global increase in 5hmC in both Hi-C A and B compartments, almost exclusively in the same 10 kb windows as loss of 5mC. This occurred despite a moderate decrease in the levels of TET1 and TET2, the major TET enzymes present in mouse ES cells [56]. The parallel increase in 5hmC and decrease in 5mC indicates that OGT normally suppresses the canonical biochemical activity of TET enzymes in mESC. A previous study using recombinant proteins suggested that OGT potentiated (rather than suppressing) TET1 activity through *O*-GlcNAcylation of its catalytic domain [25], but given the large number of potential modulators of both TET and OGT activity in cells [5–7, 12–18], the two findings are not mutually exclusive. An important question is whether TET proteins are present in heterochromatin but suppressed by OGT, or whether – as our data suggest – they are poorly localised to heterochromatin of wildtype mESC but redistribute to heterochromatin in the absence of OGT. There are many precedents for redistribution of chromatin-associated proteins in the genome: for instance, at after *Tet1* deletion in mESC, DNMT3A redistributes to genomic regions previously occupied by TET1 [60]. Likewise, the DNMT3A PWWP domain, which has greater affinity for K36me2 than for K36me3, redistributes away from intergenic and heterochromatic regions to genic regions upon depletion of the NSD1/NSD2 histone methyltransferases that generate H3K36me3 [93]. We are currently addressing these possibilities.

Cancer cells [74, 75], cells undergoing replicative senescence [76, 77] and cells from aged individuals [75, 95] all display large “partially methylated domains”, in which DNA hypomethylation occurs across megabase-sized regions corresponding to heterochromatin (reviewed in [88]). These and other pathological conditions – including cancers in general [21], BRCA1-mutant cancers [96], autoimmune/inflammatory diseases [97], neuronal dysfunction [98] and cellular senescence [77, 94] – all show the characteristic features of increased TE and satellite expression [88]. *TET2* is frequently mutated in blood cancers, but TET loss of function is likely to be a major factor in many solid cancers as well, stemming from hypoxia, decreased α-ketoglutarate levels, or increased levels of the product inhibitor succinate (reviewed in [99]). It will be important to define the functional consequences of disrupting different TET-OGT interactions not only in mESC, but also in differentiated cell types and in cancer cells. The “hypomethylating agents” 5-azacytidine and decitabine – which inhibit all DNMTs and have substantial toxicity – are used therapeutically for myelodysplastic syndrome, chronic myelomonocytic leukemia, acute myeloid leukemia and other myeloid neoplasms [100]. It would be worth exploring whether carefully titrated small molecule inhibitors of OGT [101] might be an alternative and potentially less harmful mechanism for decreasing DNA methylation in patients with these conditions.

---

**Supplementary Figure S1:**
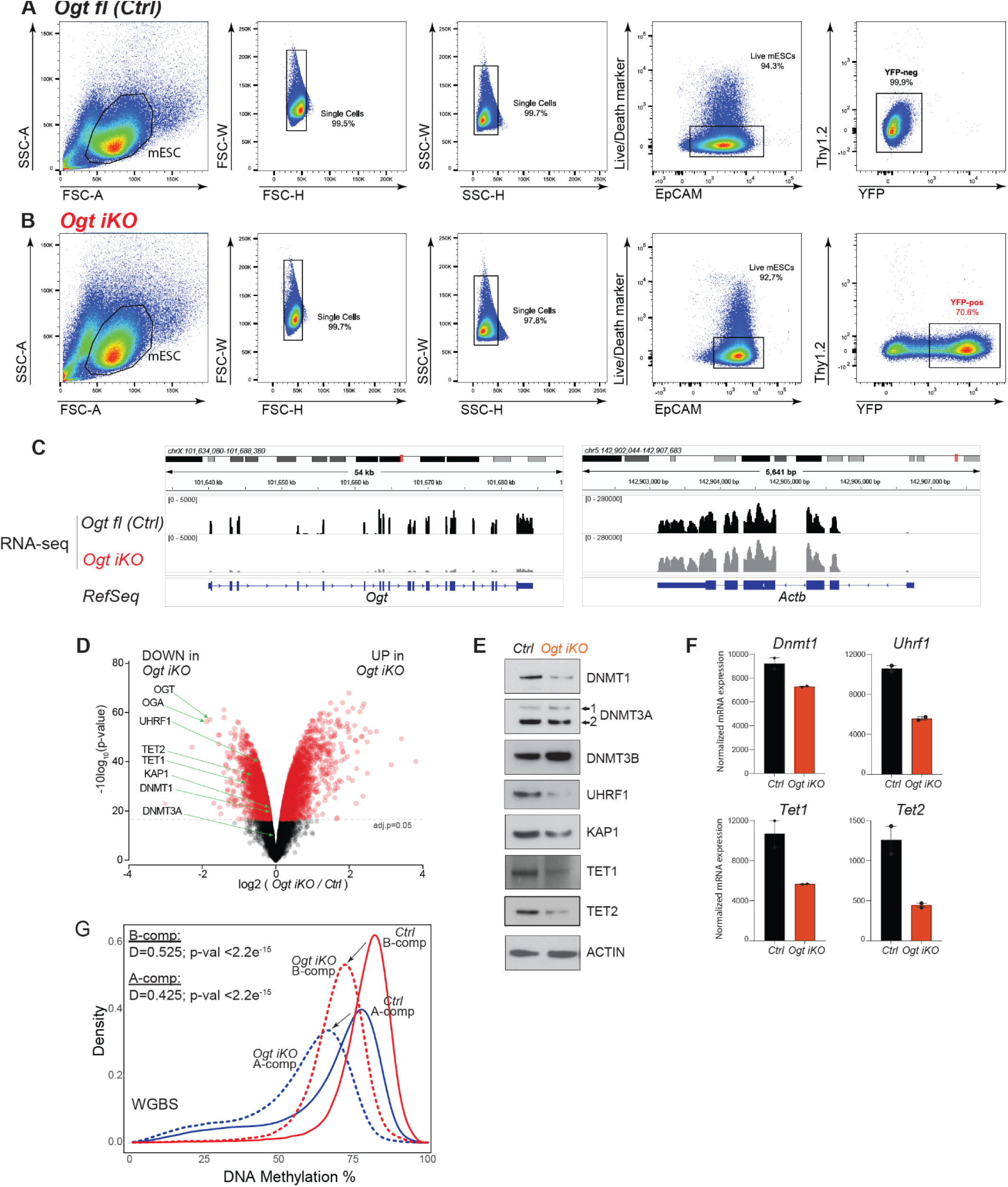
Fluorescence activated cell sorting (FACS) parameters used to isolate *Ctrl Ogt fl* and YFP^+^ *Ogt iKO* mESC. **A**-**B**, Flow cytometry gating scheme for analysis of sorted *Ctrl Ogt fl (***A**) and *Ogt iKO* (**B**) mESC. The sorting strategy includes successive staining for Live/Dead, Thy1.2 (to eliminate MEFs – mouse embryonic fibroblasts in the feeder layer) and EpCAM (to detect mESC). After 6 days of exposure to 4-OHT, *Ogt fl* cells yielded >70% YFP^+^ *Ogt iKO* mESC in which Cre had been activated; these cells, as well as YFP-negative cells from *Ctrl Ogt fl* mESC that had not been exposed to 4-OHT, were sorted and used for further analyses. **C**, *Left*, Genome browser view of RNA-seq data from the *Ogt* locus showing the considerable decrease of *Ogt* mRNA in *Ctrl Ogt fl* compared to *Ogt iKO* mESC. The *ActB* locus was used as control (*right*). **D-E**, Changes in the levels of selected proteins in *Ogt iKO* compared to *Ctrl Ogt fl* mESC shown by **D**, volcano plot of mass spectrometry data [21] and **E**, western blotting (with actin used as loading control). Note the substantial decrease in DNMT1, UHRF1 and TET2 protein levels by western blotting, and the more moderate decrease in DNMT3A1/2, KAP1 and TET1. DNMT3B protein levels show a substantial increase. **F**, normalized RNA expression of *Dnmt1, Uhrf1, Tet1* and *Tet2* in *Ogt iKO* compared to *Ctrl Ogt fl* mESC, assessed by RNA-seq. **G**, Density distribution plot of average 5mC+5hmC (WGBS) values in 10 kb windows across the genome, showing loss of 5mC+5hmC in both euchromatin (*blue*) and heterochromatin (*red*) of *Ogt iKO* (*dashed line*) compared to *Ctrl* (*solid line*) mESC.

**Supplementary Figure S2:**
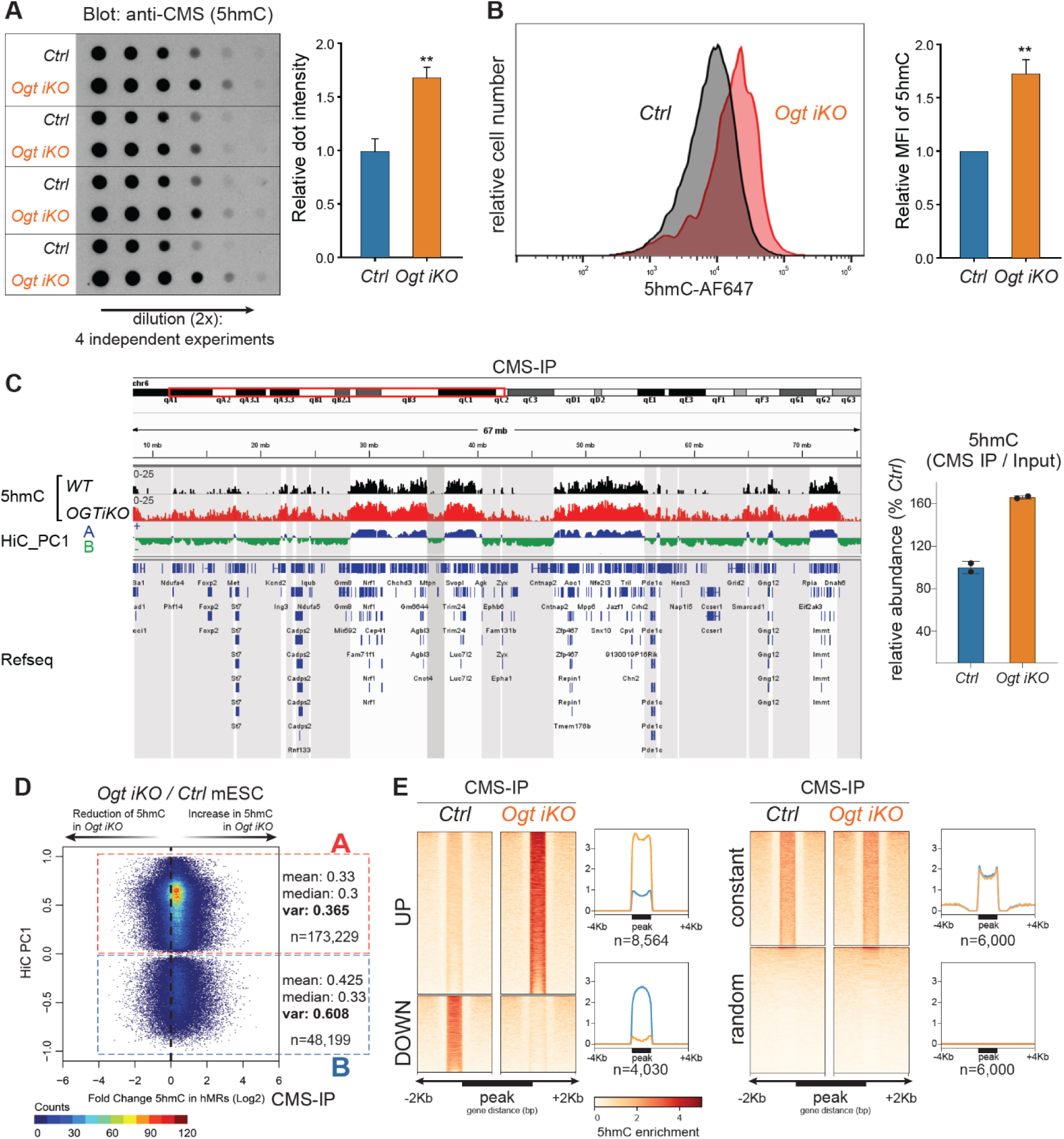
Increased 5hmC levels in *Ogt iKO* compared to *Ctrl* mESC, assessed using DNA dot blot, flow cytometry and CMS-IP. **A**, *Left*, DNA dot blot assay using antibodies to the 5hmC derivative cytosine-5-methylenesulfonate (CMS) to assess global 5hmC levels in *Ogt iKO* compared to *Ctrl Ogt fl* mESC. Data are representative of four independent experiments. *Right*, Quantification of relative dot intensity. There is a very reproducible ∼1.6-fold increase in 5hmC in *Ogt iKO* compared to *Ctrl Ogt fl* mESC, using all three assays described below. **B**, *Left*, 5hmC levels in *Ogt iKO* and *Ctrl* mESC assessed by flow cytometry with an anti-5hmC antibody. A representative histogram is shown. *Right*, Quantification of relative mean fluorescence intensity (MFI) of 5hmC staining in 3 independent experiments in *Ogt iKO* compared to *Ctrl Ogt fl* mESC. **C**-**E**, 5hmC genomic distribution assessed by CMS-IP in *Ogt iKO* and *Ogt fl (Ctrl)* mESC. Spike-in controls were used for internal normalization. **C**, *Left*, Genome browser view of 5hmC levels in *Ogt iKO* and *Ctrl Ogt fl* mESC. *Right*, Quantification of normalized reads in CMS-IP peaks. **D**, Dot plot showing increased 5hmC (quantified as normalized CMS-IP reads) in 5hmC-rich regions in *Ctrl Ogt fl* and *Ogt iKO*, plotted in euchromatin (Hi-C A) and heterochromatin (Hi-C B) compartments. Each dot represents a hydroxymethylated region (hmRs). **E**, Heat maps of differentially hydroxymethylated regions (DhmRs) in *Ogt iKO* compared to *Ctrl Ogt fl* mESC.

**Supplementary Figure S3:**
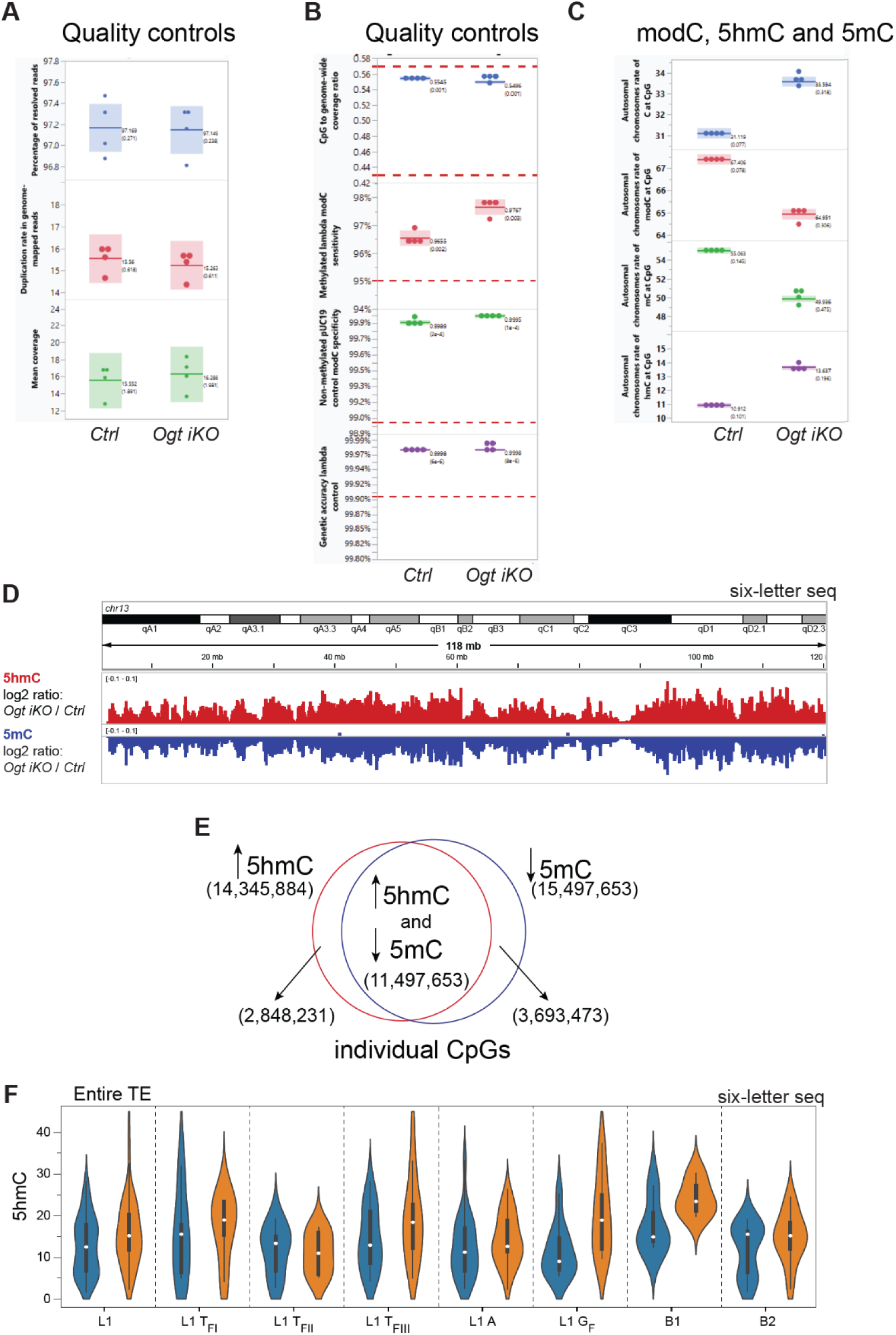
Quality controls and additional data for six-letter sequencing in *Ogt iKO* and *Ctrl Ogt fl* mESC. **A**, Average percentage of resolved reads. *Top*, reads resulting from computationally resolving read 1 (original) and read 2 (copy) strands of a six-letter sequencing construct from a paired-end illumina sequencing run. Read 1 and read 2 resolves to single reads containing bases in one of six states (A, C, G, T, 5mC or 5hmC) [50]. *Middle*, Duplication of genome-mapped resolved reads. *Bottom*, Mean coverage of all genome-mapped resolved reads. **B**, CpG to genome-wide coverage. *Top*, Ratio of mean coverage at CpGs to mean coverage genome-wide. Note: CpGs are stranded with coverage compared to overall coverage on both strands (therefore ideal coverage is 0.5). *Second from top*, Fully methylated lambda genome spike-in control mC sensitivity). *Third from top*, Fully unmethylated pUC19 genome spike-in control C specificity. *Bottom*, Genetic accuracy of lambda genome spike-in control. **C**, *Top*, Autosomal chromosome rate of C at CpG sites. *Second from top*, modC at CpG sites. *Third from top*, 5mC at CpG sites. *Bottom*, 5hmC at CpG sites. **D**-**E**, The increase of 5hmC and decrease of 5mC in *Ogt iKO* compared to *Ctrl Ogt fl* mESC occurs in the same genomic regions. **D**, Genome browser view showing the reciprocal gain of 5hmC (*red*) and loss of 5mC (*blue*) in *Ogt iKO* compared to *Ctrl Ogt fl* mESC. Plotted are log2 ratios (*Ogt iKO*/ *Ctrl Ogt fl*) for 5hmC (*top*) and 5mC (*bottom*). **E**, Venn diagram showing the overlap between CpGs gaining 5hmC (*blue*) and those losing 5mC (*red*) in *Ogt iKO* compared to *Ctrl Ogt fl* mESC. **F**, 5hmC levels averaged across the entire transcribed units of individually mapped TEs belonging to the indicated TE subfamilies and showing increased expression in *Ogt iKO* compared to *Ctrl Ogt fl* mESC. Like expressed genes which have high 5hmC in their gene bodies, expressed TEs are enriched for 5hmC in their transcribed regions as well as in their regulatory regions (Fig. 3F).

**Supplementary Figure S4:**
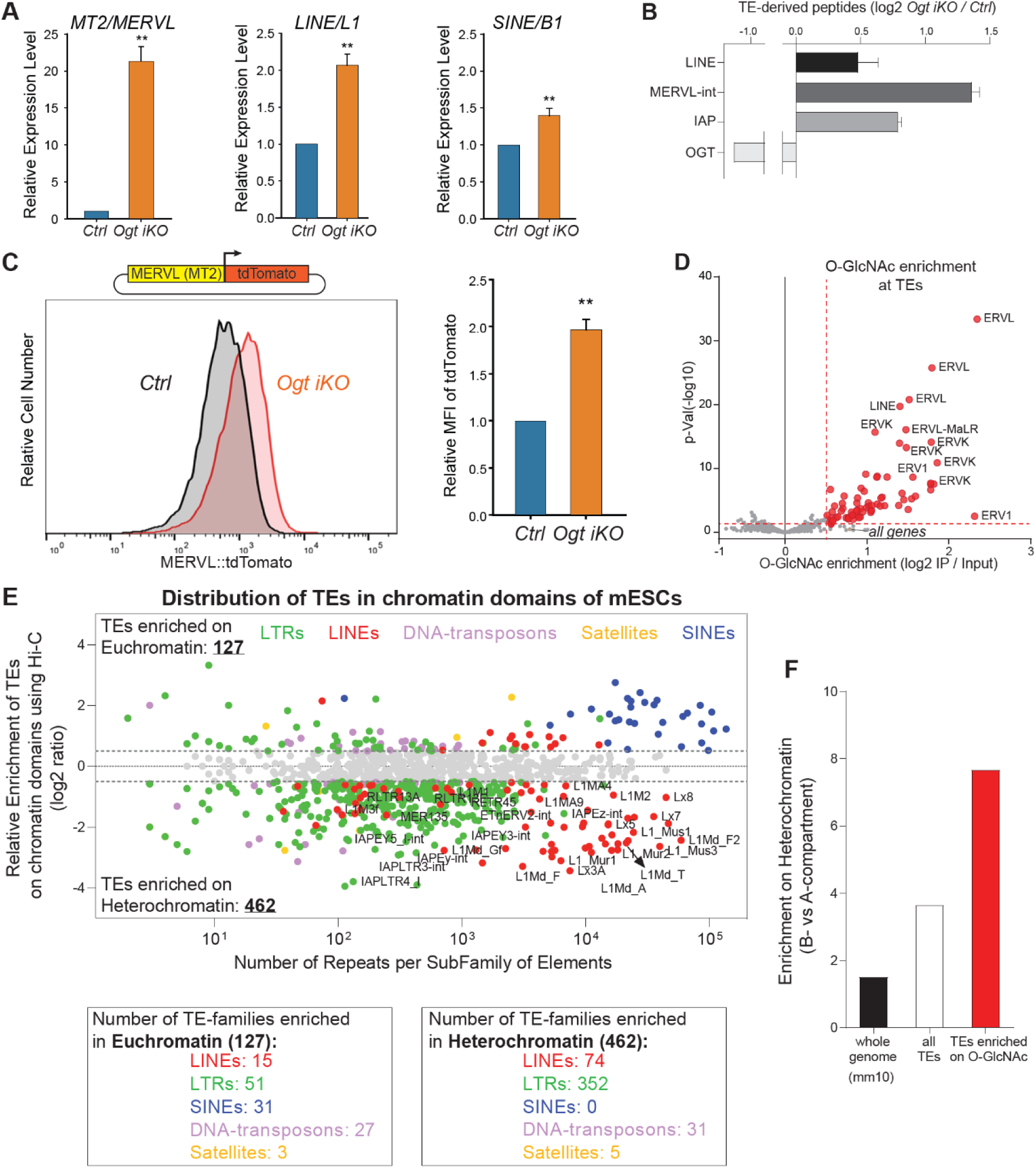
TEs regulated by OGT are preferentially enriched in heterochromatic regions. **A**, Increased expression of MT2/MERVL and LINE1/L1 in *Ogt iKO* compared to *Ctrl Ogt fl* mESC, assessed by RT-qPCR. **B**, Increased detection of peptides from LINE L1, MERVL-int and IAP elements in *Ogt iKO* compared to *Ctrl Ogt fl* mESC. Mass spectrometry samples of whole-cell lysates were analyzed against a reference dataset of proteins encoded in transposable elements (*see Methods*). **C**, Increased tdTomato fluorescence in *Ogt iKO* mESC transfected with a tdTomato reporter driven by the MERVL-MT2 LTR. *Upper left,* schematic of the *MERVL::tdTomato* reporter*; lower left*, representative flow cytometry histograms, *Right*, quantification of mean fluorescence intensity (MFI) in three independent experiments. The MERVL LTR reports on the 2C-like state in mESC. **D**, *O*-GlcNAc enrichment at chromatin-associated proteins located within TEs, based on *O*-GlcNAc ChIP-seq [33]. **E**, Presence of TE subfamilies in euchromatin or heterochromatin domains in *WT* mESC. *Top*, For each subfamily of TEs (represented as a single dot), we counted the number of individual TE copies contained in euchromatin (positive Hi-C PC1 values) or in heterochromatin (negative Hi-C PC1 values). The log2 ratio of the number of TE copies contained in euchromatin versus heterochromatin are plotted on the y-axis while the total number of copies found in the genome (mm10) are shown on the x-axis. Enriched TEs, log2 ratio > or < +/-0.5. Color codes, TE classification by subfamilies (shown at top). Gray dots, TE-subfamilies with no preferential presence in euchromatin or heterochromatin. *Bottom*, total number of TE subfamilies enriched in euchromatin or heterochromatin by classes of TEs (LINEs, LTRs, SINEs, DNA-transposons, and Satellites). **F**, TEs containing high levels of the *O*-GlcNAc modification are enriched in heterochromatin. Ratios of TEs in heterochromatin versus euchromatin are shown for the whole mouse genome (*black bar*), for all TEs (*white bar*) and for those TEs that are enriched in O-GlcNAc (as shown in Fig. 4A and **Suppl. Fig. 5**; *purple bars*).

**Supplementary Figure S5:**
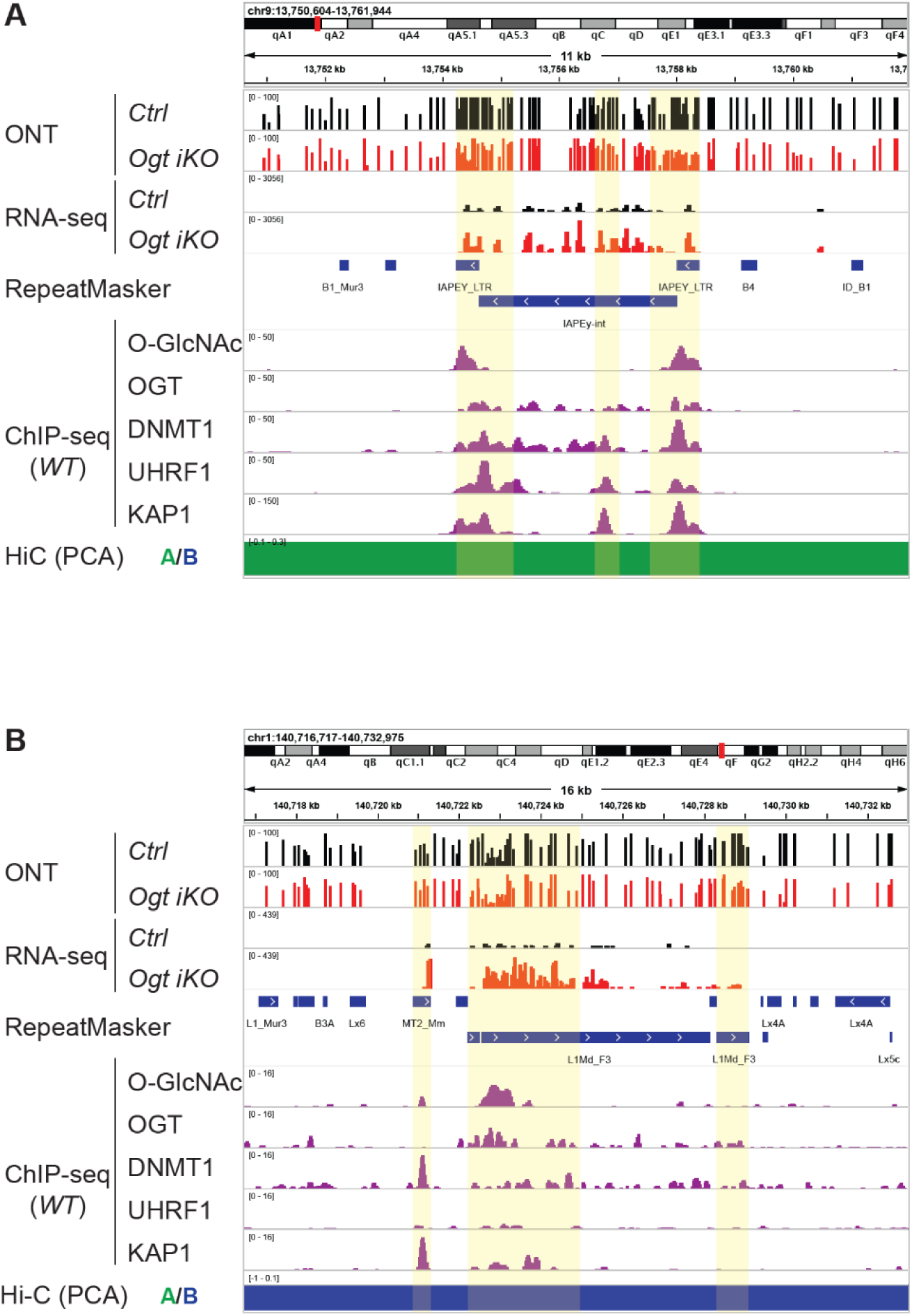
Activation of individual L1 and IAP elements is accompanied by loss of DNA methylation at their regulatory regions in *Ogt iKO* mESC. In all figures, the top four tracks show ONT-seq and RNA-Seq data from *Ctrl* and *Ogt iKO* mESC, followed by HiC data (PC1) indicating the presence of these regions at euchromatin (green) or heterochromatin (blue). *Bottom tracks*, ChIP-Seq for *O*-GlcNAc, OGT, DNMT1, UHRF1 and KAP1 in WT mESC [22, 33, 64–67, 86]. **A**, Genome browser view of a uniquely mapped IAP element in euchromatin. Note the loss of DNA modification (ONT sequencing) at certain CpGs (*ONT, top tracks*), the increased expression of the IAP internal region as well from the flanking LTRs in *Ogt iKO* compared to *Ctrl Ogt fl* mESC (RNA-seq tracks), and the presence of the *O*-GlcNAc modification on chromatin-associated proteins occupying the LTR regulatory regions, based on *O*-GlcNAc ChIP-seq in mESC [33]. **B**, Genome browser view of a uniquely mapped L1Md_F3 element in heterochromatin. Note the loss of DNA modification (ONT sequencing) at certain CpGs (*ONT, top tracks*), the increased expression of the L1 as well as the nearby MT2 element in *Ogt iKO* compared to *Ctrl Ogt fl* mESC (RNA-seq tracks), and the presence of the *O*-GlcNAc modification on chromatin-associated proteins occupying the 5’UTR regulatory region, based on *O*-GlcNAc ChIP-seq in mESC [33].

**Supplementary Figure S6:**
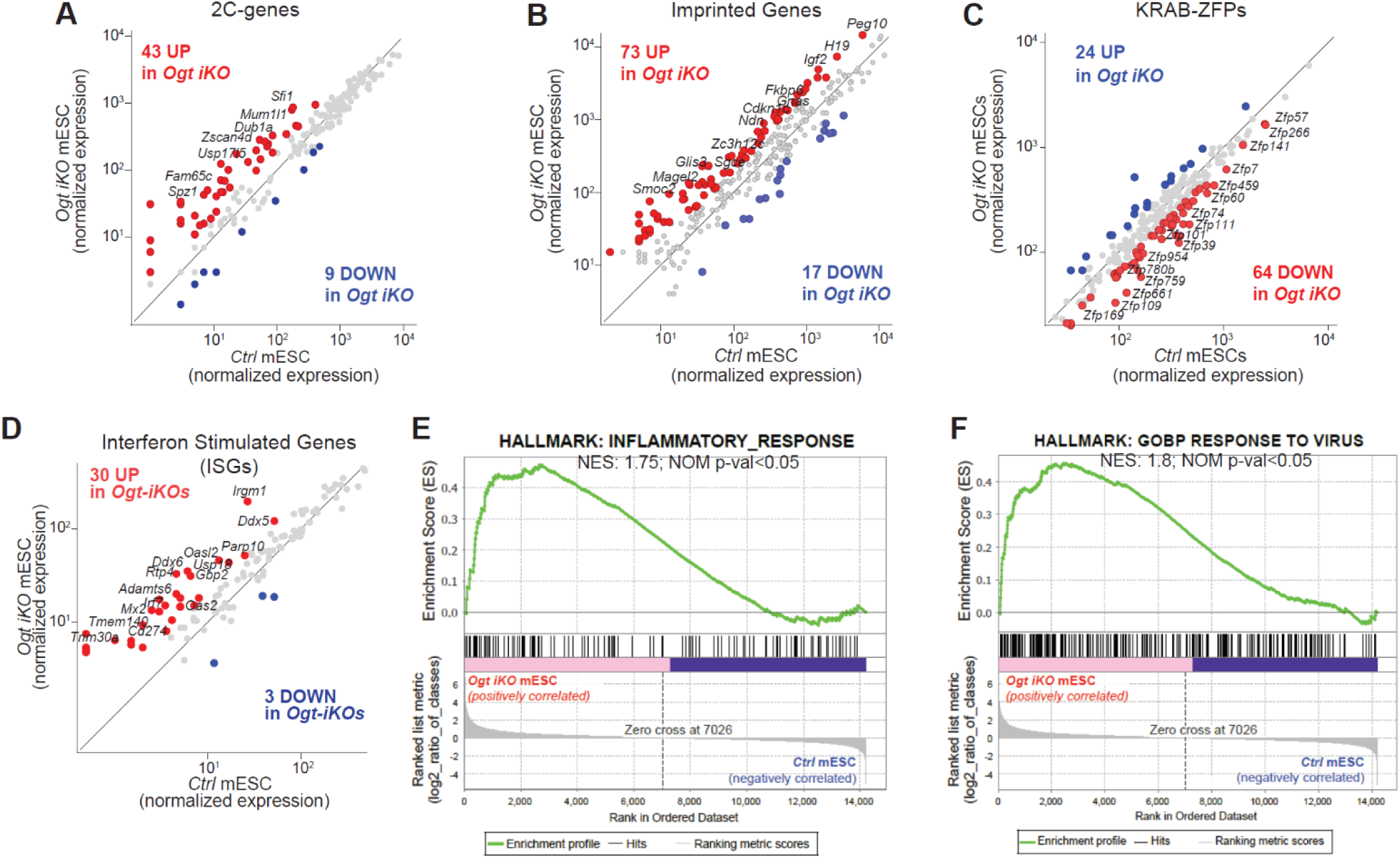
Transcriptional signatures of *Ogt iKO* mESC. **A-D**, Scatter plots comparing expression of **A**, genes related to the 2C-like state; **B**, imprinted genes; **C**, Krüppel-associated box domain (KRAB) zinc finger (ZNF) genes; and **D**, Interferon-stimulated genes (ISGs) in *Ogt iKO* versus *Ctrl Ogt fl* mESC. **E-F**, Gene Set Enrichment Analysis (GSEA) comparing *Ogt iKO* versus *Ogt fl (Ctrl)* mESC. Enriched pathways in *Ogt iKO* mESC include **E**, Inflammatory response and **F**, Response to virus.

**Supplementary Figure S7:**
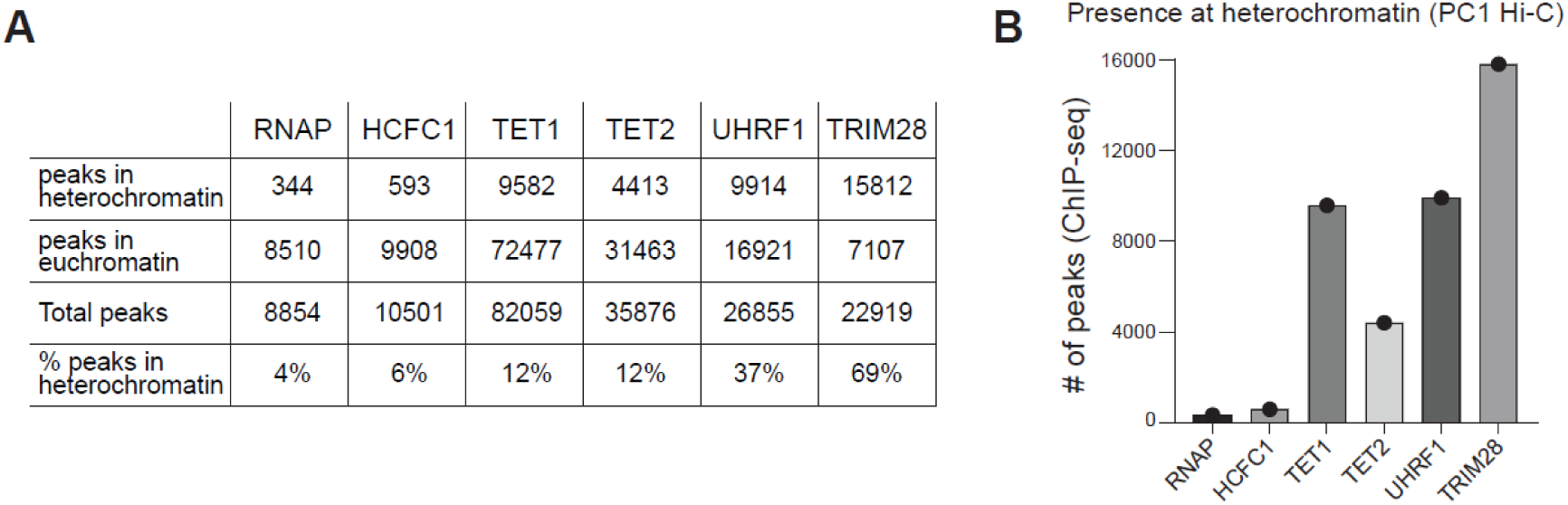
TET proteins are preferentially enriched in euchromatin and bind at some heterochromatin regions in mESC. **A**, Table quantifying numbers of TET1 and TET2 ChIP-seq [64, 65] peaks, as well as ChIP-seq peaks for other chromatin proteins, in euchromatin or heterochromatin (Hi-C PC1 values). A minority of RNA polymerase II (RNAP), HCFC1, TET1 and TET2 ChIP-seq peaks (4%, 6%, 12% and 12% respectively) are located in heterochromatin, compared to UHRF1 (37%) and TRIM28 (69%). **B**, Graph showing the total number of ChIP-seq peaks located in heterochromatin for TET1, TET2 and the chromatin proteins mentioned above. The number of TET1 ChIP-seq peaks in heterochromatin are similar to the number for UHRF1, and much greater than the numbers for RNAP and HCFC1.

**Supplementary Figure S8:**
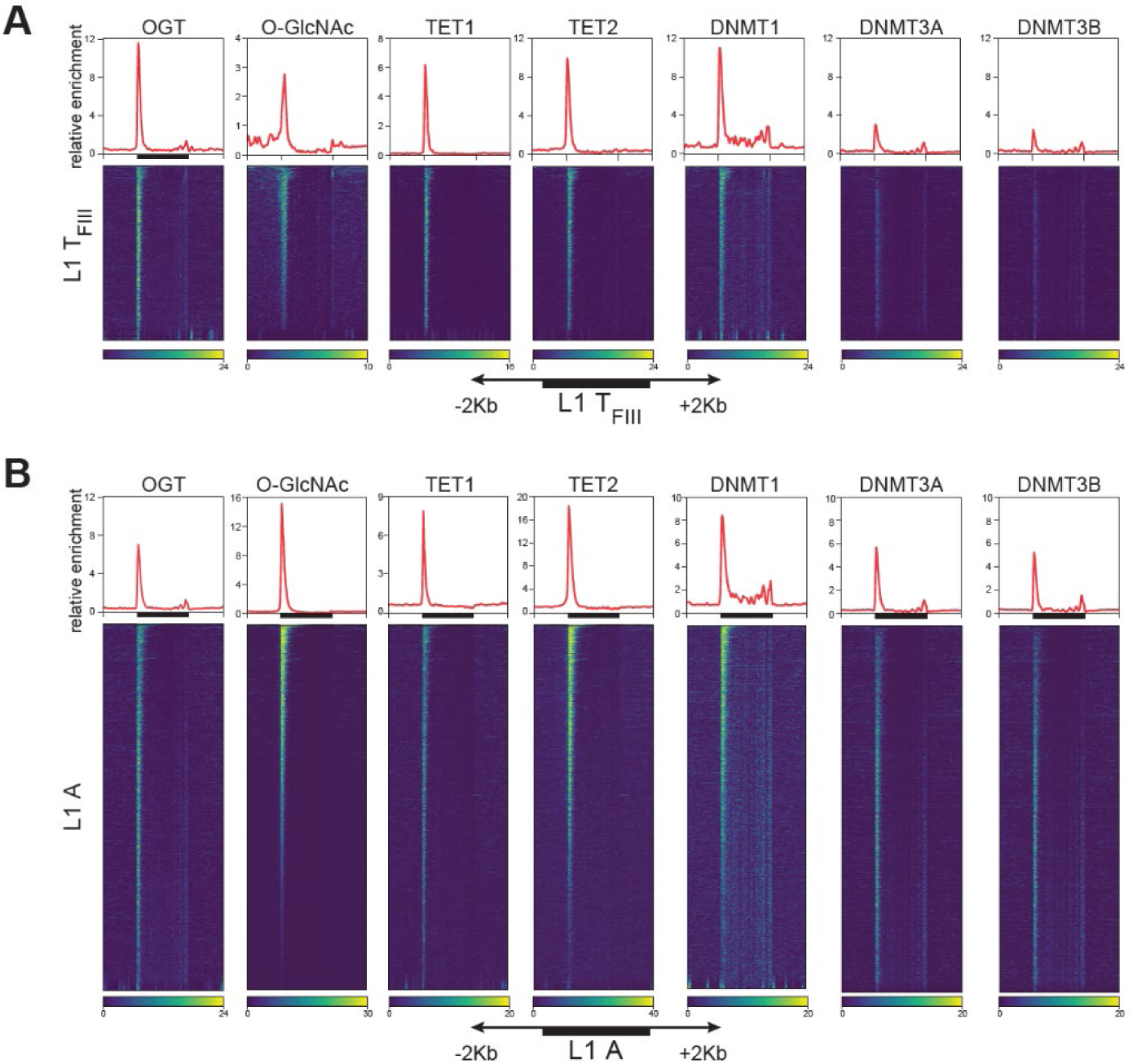
Enrichment of OGT, *O*-GlcNAc, TET1, TET2 and all three DNMTs at the 5’ regulatory regions of the relatively young family of LINE elements, L1 T_FIII_ and L1 A in WT mESC. Heatmaps were generated using ChIP-seq datasets [22, 33, 64–67, 86] for the TE subfamilies LINE elements, L1 T_FIII_ (**A**) and L1 A (**B**) in WT mESC.

## MATERIALS AND METHODS

### Generation and Culture of *Ogt iKO* mouse Embryonic Stem Cells (mESC)

The *Ogt-inducible knockout (Ogt-iKO; Ubc-Ogt^floxed^; Cre–ERT2^+/KI^; Rosa26-LSLYFP^+/KI^)* mESC was previously generated by our group [21]. *Ogt* deletion was induced by exposing *Ogt iKO* mESC to 1 μM 4-hydroxytamoxifen (4-OHT) (TOCRIS, 3412). At the time points used in this study (6 days after 4-OHT treatment), *Ogt iKO* mESC have undergone G1 arrest but are still fully viable [21]. mESC were cultured over mitomycin C-treated mouse embryonic fibroblasts (MEFs) as feeder cells and using knockout DMEM medium (ThermoFisher Scientific, 10829018) supplemented with LIF, 15% KOSR (KnockOut Serum Replacement; ThermoFisher Scientific, 10828028), GlutaMAX^TM^ (Gibco, 35050061), 1x MEM nonessential amino acids (Gibco, 11140-035) and 50 μM β-mercaptoethanol (Gibco, 21985023). To generate *Ogt iKO* mESC carrying the 2C-reporter we used the MERVL::tdTomato reporter [51] that was kindly shared from Samuel Pfaff (Addgene #40281). Reporter stable cell line was generated by transfection (Lipofectamine™ 2000, thermofisher) and drug selection (Hygromycin, Thermofisher).

### Cell sorting and sample generation

*Ogt iKO* mESC were cultured during 6-days in presence of 4-OHT to generate OGT-depleted mESC that efficiently express YFP+ (>85% of the entire population; Supp Fig1) that results from a successful Cre-ERT2-mediated recombination, while control mESC (*Ctrl*) were maintained without 4-OHT. In order to ensure a high purity of our samples, we isolate specifically the YFP^+^ population (in the case of 4-OHT-treated *Ogt iKO* mESC) and also to eliminate the presence of mitomycin-C-treated MEFs (used as feeders during our mESC cultures) we performed FACS (Fluorescence-Activated Cell Sorting) using a FACSAria cell sorter (BD Biosciences). The dissociated mESC were first labeled with the eBioscience^TM^ Fixable Viability Dye eFluor^TM^ 780 (ThermoFisher, 65-0865-14) to select viable cells and then stained for surface markers with indicated fluorochrome-conjugated antibodies diluted in FACS buffer (PBS 1X, 0.5% BSA, 0.01% NaN_3_): CD90.2 (Biolegend, 140319) to positively distinguish the mESC and CD326 (Biolegend, 118227) to negatively exclude MEFs.

### RNA-Seq

Total RNA was isolated from *Ogt iKO* and *Ctrl* mESC samples using RNeasy plus mini or micro kit (Qiagen). RNA-sequencing libraries were prepared using Truseq stranded mRNA kit (Illumina) according to the manufacture’s protocol. All libraries were assessed using Qubit™ RNA HS Assay Kit (ThermoFisher, Q32855) and TapeStation High Sensitivity RNA ScreenTape Analysis (Agilent, 5067-5579). Libraries were pooled in equal quantity and sequenced using Illumina HiSeq 2500 (Illumina) to produce 20 million paired-end reads (50×50 bases) per each RNA-Seq library. RNA-seq data were deposited under the GEO accession number GSE252760.

### Whole genome bisulfite sequencing (WGBS)

Genomic DNA was isolated using PureLink Genomic DNA Mini Kit (Thermo Fisher Scientific, K182001) or FlexiGene DNA Kit (Qiagen, 51206). Unmethylated lambda DNA (Promega, D1521) was spiked into the genomic DNA samples at a ratio of 1:200 to monitor the bisulfite conversion efficiency. The samples were then sheared using Covaris S2, purified with Ampure XP beads (Beckman Coulter) and processed with NEBNext End Repair and dA-Tailing Modules (NEB), and ligated to methylated Illumina Adaptors using NEBNext Quick Ligation Module (NEB). DNA with ligated adaptors was then treated with sodium bisulfite (MethylCode, Thermo Fisher Scientific) for 4hrs and amplified with index primers using KAPA HiFi HotStart Uracil+ Ready Mix (KAPA Biosystems). The samples after PCR amplification were then purified using Ampure XP Beads (Beckman Coulter) and sequenced as 125- or 250-base-pair paired end reads using Illumina Hiseq 2500 (Illumina). WGBS data were deposited under the GEO accession number GSE252760.

### Oxford Nanopore Technologies sequencing (ONT-seq)

After cell sorting, High Molecular Weight genomic DNA was isolated from *Ogt-iKO* and *Ctrl* mESCs using Nanobind CBB kit (Circulomics, NB-900-001-01) according to the manufacturer’s instructions. DNA libraries were prepared at the Kinghorn Centre for Clinical Genomics (KCCG, Australia) using 3 μg input DNA, without shearing, and an SQK-LSK110 ligation sequencing kit. Libraries were each sequenced separately on a PromethION (Oxford Nanopore Technologies) flow cell (FLO-PRO002, R9.4.1 chemistry). Bases were called with guppy 5.0.13 (Oxford Nanopore Technologies). ONT sequencing data were deposited in the European Nucleotide Archive (ENA) under identifier PRJEB67735.

### Cytosine-5-methylenesulfonate immunoprecipitation (CMS-IP)

CMS-IP to detect 5hmC was performed as described before [9, 102, 103], using 3 ug of genomic DNA as starting material. After library preparation, Next generation sequencing was performed to generate 30 million reads of IP samples as well as from Input samples. Equal amounts of a PCR amplicon generated to contain 5hmC (5hmC spike-in) were added to all samples for internal calibration. CMS-IP data were deposited under the GEO accession number GSE252760.

### Simultaneous detection of 5hmC and 5mC by six-letter method

The six-letter method was performed as described before [50], using 100 ng of genomic DNA as starting material from *Ogt iKO* cells sorted to be YFP^+^ after tamoxifen treatment, and control YFP-negative cells sorted without prior treatment with tamoxifen. six-letter sequencing data have been deposited in the NCBI Gene Expression Omnibus and are accessible through the GEO Series accession number GSE247534.

### Flow cytometry

mESC were trypsinized into single cells and used for staining or cell sorting. For 5hmC staining, the genomic DNA was denatured by treating the fixed mESC with 2 N HCl for 30 min. Following the removal of HCl, the pH was neutralized with 100 mM Tris-HCl (pH 8.5) for 10 min. at room temperature. For staining, we used Active Motif #39791 as primary antibody (anti-5hmC) and Thermofisher #A32795 as secondary antibody (IgG).

### Quantitative real-time PCR (qRT-PCR)

Quantitative real-time PCR was performed using Universal SYBR Green Master Mix (Roche) and analyzed using a Step One Plus real-time PCR system (Applied Biosystems) according to the manufacturer’s instructions and the data were normalized for *Gapdh* expression. The primers used for qRT-PCR are listed in Table.

### Bioinformatic Analyses

For *RNA-Seq* data, reads were aligned using STAR [104] with the parameters --outFilterMultimapNmax 1, -- outSAMtype BAM SortedByCoordinate –sjdbOverhang 100 to analyze gene expression. HT-Seq [105] was used to quantify the gene expression levels using the options htseq-count -s yes, -r pos, -a 10. Normalization and differential expression analyses were performed using DESEq2 [106], with the parameters fitType parametric, alpha 0.05 and using the Benjamini & Hochberg method. For visualization of the data, we generated tracks using Deeptools [107], with the option bamCoverage. All related plots were made using R-Studio [108] and Integrative Genome Viewer (IGV) [109]. To measure the expression of Transposable Elements (TEs), we used the TE-transcript package [110] with the option -multi to use ambiguously mapped reads and then perform the differential expression (DE) analysis with DESEQ2 (FDR cutoff of p<0.05). Genes or TEs with less than 10 reads total were pre-filtered in all comparisons as an initial step. In all our analysis, we used the mm10 genome version and for analysis of TEs, we specifically used the mm10 repeatMasker version from TE-transcript [110]. For detection of individual TEs we used the STAR aligner with the parameter --outFilterMultimapNmax 1 to then count the amount of uniquely aligned reads using htseq-count against the customized mm10 version from Hammell lab as genome reference.

For *ChIP-Seq* datasets, we used Bowtie [110] for alignments and Deeptools [107] with the option bamCoverage for generation of the genome tracks. Heatmaps and profiles were generated using computeMatrix with the option scale-regions and plotHeatmap, both from Deeptools.

A *Hi-C* dataset from *WT* mESC [112] was used for genome compartmentalization. We estimated the relationship between the number of TE-copies contained in euchromatin (positive PC1 values) versus heterochromatin (negative PC1 values) using a logarithmic transformation, using the formula: log 2 ((number of elements in Euchromatin)/(number of elements in Heterochromatin)); the resulting values are plotted in the y-axis while the total number of copies found in the genome (mm10) are shown in the x-axis. Positive values in the y-axis (upper side of the plot) show TE-subfamilies which copies are enriched in Euchromatin while negative values (bottom side of the plot) show TEs-subfamilies which copies are enriched in heterochromatin. Enriched TEs were defined as those having log2 ratio > or < +/-0.5.

For *Whole Genome Bisulfite sequencing (WGBS)* samples we used Bismark [113] and bedGraphToBigWig for tracks generation. The median coverage for each sample was ∼20X.

For WGBS, as well as for ONT and six-letter-seq data, we used CpGs covered by 5 or more reads for the analyses. Additionally, for analysis using windows of 10 kilobases (10 kb), we only included those regions containing more than 5 CpGs.

For *ONT data*, reference TE methylation was assessed for mESC samples aggregated by condition using Methylartist [114] version 1.2.482. Briefly, CpG methylation calls were generated from ONT reads using nanopolish version 0.13.2118. Using Methylartist commands db-nanopolish, segmeth and segplot with default parameters, methylation statistics were generated for the genome divided into 10kbp bins, protein-coding gene promoters defined the Eukaryotic Promoter Database (−1000bp,+500 bp)119, and reference TEs defined by RepeatMasker coordinates deposited in the TE-transcript package [110]. The median coverage for each sample was ∼10X.

For *six-letter-seq* data was processed as before [50] and the median coverage for each sample was ∼15X.

For CMS-IP, IP and Input samples were aligned against the mm10 genome reference using Bismark [113] version v0.15.0. Reads aligning to a PCR amplicon generated to contain 5hmC (5hmC spike-in) that was equally present across samples were used for internal calibration, as described before [103]. Peak calling was performed with MACS2, using Input DNA to compare against the CMS-IP samples. Peaks from different replicates were merged using bedtools.

### Western Blots

Whole-cell extracts (WCE) were prepared by incubating mESC with radioimmunoprecipitation assay (RIPA) buffer (Thermo Fisher, 89900) supplemented with Benzonase (Sigma-Aldrich, E1014-25KU) and Halt™ Protease/Phosphatase Inhibitor Cocktail (ThermoFisher, 78441) on ice for 1 hour. Proteins from WCEs were resolved using NuPAGE 4-12% bis-tris gel (Thermo Fisher Scientific, NP0321BOX) and transferred onto polyvinylidene difluoride (PVDF) membranes using Wet/Tank Blotting Systems (Bio-Rad). PVDF membranes were blocked with 5% nonfat milk in PBST (PBS 1X, 0.05% Tween-20) and incubated with the indicated primary antibodies, followed by secondary antibodies conjugated with horseradish peroxidase (HRP) (Cell Signaling, 7076V), all of which were diluted 5% nonfat milk in PBST. Signal was detected with Femto Supersignal ECL substrate (ThermoFisher, 34096) and X-ray films (ThermoFisher, 34090).

### Generation of endogenously flagged OGT (*Ogt-flag*) mESC and sample generation

Stable epitope-tagged OGT mouse embryonic stem cell lines were generated using CRISPR/Cas9 homology directed repair (HDR) to knock in the epitopes flag or streptavidin into the endogenous *Ogt* locus. Nucleofection with plasmid containing the repair template, an *in vitro* transcribed sgRNA (EnGen, NEB), and recombinant CAS9 was done using Lonza’s P1 kit, following manufacturer’s recommendations. Antibiotic selection was performed by culturing the transfected mESC with 1 um puromycin (Thermo Fisher, A1113803) for 2 weeks. Stable *Ogt*-edited mESC were confirmed by detecting the expression of the OGT-tagged versions by western blot.

### Mass Spectrometry

For sample generation, approximately 30 million epitope-tagged OGT mESC stable cell lines were trypsinized and washed twice with cold 1x PBS before placing on ice. Subcellular fractionation into nuclear and cytoplasmic fractions was done using Thermo Fisher NE-PER Nuclear and Cytoplasmic Extraction kit (78833), supplemented with protease inhibitor cocktail (Thermo 78429), as described in the manufacturer’s protocol. Protein quantification was done using Bradford colorimetric assay (BioRad 5000006). Anti-FLAG M2 magnetic beads (Sigma M8823) were washed with RIPA buffer before adding 1ul beads/1mg protein to protein samples and undergoing continuous inversion at 4°C overnight. Proteins were processed and analyzed as previously described with minor modifications [115]. Samples were analyzed on an Orbitrap Eclipse mass spectrometer (Thermo Fisher Scientific).

### Mass Spectrometry data analysis

Raw data were processed using MaxQuant software [116] using the default settings and searched against the mouse UniProt database (12/2017) with common contaminant entries. The settings used for MaxQuant analysis were: enzymes set as LysC/P and Trypsin/P, with maximum of 2 missed cleavages; fixed modification was Carbamidomethyl (Cys); variable modifications were Acetyl (Protein N-term) and Oxidation (Met); label-free quantification was used with a minimum ratio count of 2 and classic normalization; False Discovery Rate for both protein and peptide identification was 0.01. The ‘Match between runs” feature was enabled. The proteinGroups file from the MaxQuant analysis was used as the input for all downstream analyses. Gene names that were not automatically assigned by MaxQuant were manually added from their Entrez protein IDs. Of the 2,617 entries in the proteinGroups file, we filtered out 33 entries identified as contaminants. Each of the three replicates within a condition (flag-tagged or strep-tagged/control) were annotated. LFQ intensities were log2 transformed and proteins with less than 2 LFQ intensity values across both conditions were filtered out. Missing values were imputed with the minimum detected intensity. A two-tailed t-test with a p-value cutoff of 0.05 and FDR of 0.01 was conducted alongside the calculation of log2 fold changes for construction of the volcano plot.

## ACKNOLEDGMENTS

We thank C. Kim and S Alarcon and colleagues of the La Jolla Institute (LJI) Flow Cytometry and Next-Generation Sequencing Facilities for help with cell sorting and next-generation sequencing, respectively. The FACSAria II Cell Sorter (S10 RR027366), the NovaSeq 6000 (S10OD025052) and the HiSeq 2500 (S10 OD018499) were acquired through the Shared Instrumentation Grant (SIG) Program. ONT sequencing was done using a PromethION at the Garvan Sequencing Platform in Sydney. This research was supported by NIH grants R35 CA210043 to A.R., NIH NIGMS R35GM147554 to S.A.M. and NHMRC Investigator Grant (GNT1173711) and the Mater Foundation to G.J.F. H.S. was supported by the Pew Latin-American Fellows Program from The Pew Charitable Trusts, and by a Fellowship from the California Institute for Regenerative Medicine, X.L. was supported by a Fellowship from the California Institute for Regenerative Medicine, M.B. was supported by the UCSD Graduate Training Program in Cellular and Molecular Pharmacology through the institutional training grant NIH NIGMS T32 GM007752.

## AUTHOR CONTRIBUTIONS

H. S. and X.L. performed the experiments using the Ogt iKO mESC, H.S., J.C.A. and L.J.A.V. performed the bioinformatics analyses, N.J. prepared the ONT libraries and did the ONT sequencing, G.J.F. and H.S performed and interpreted the ONT methylation data related to transposable elements, X.Y. and C.B. helped to perform experiments, P.K. generated the 3xflag-Ogt mESC and M.B. performed the IP-mass spectrometry experiments, S.A.M. supervised the mass spectrometry experiments, S.A.M, M.B and C.M. analyzed the mass spectrometry data, the biomodal team prepared and processed the six-letter sequencing libraries, H.S. and A.R. wrote the manuscript. All authors were involved in reviewing and editing the manuscript.

